# Genome-wide diversity of coconut from northern South America uncovers genotypes present in Colombia and strong population structure

**DOI:** 10.1101/825398

**Authors:** Jorge Mario Muñoz-Pérez, Gloria Patricia Cañas, Lorena López, Tatiana Arias

## Abstract

Coconut palms (*Cocos nucifera*) are a combination of wild admixed populations and perennial crops with a worldwide distribution. Here we develop single nucleotide polymorphisms (SNPs) along the coconut genome based on Genotyping by Sequencing (GBS) for at least four different commercially important and widely cultivated coconut varieties and hybrids growing in northern South America. We present a comprehensive catalog of approximately 27K SNPs to conduct genetic diversity, population structure and linkage disequilibrium analysis. A relatively fast LD decay for the Atlantic accessions within ~250Kb was observed in comparison to the Pacific accessions ~ 1500 Kb.

The complete SNPs sampling showed a strong population structure at K = 2, separating accessions from the Pacific and Atlantic coasts as it has been found in previous studies. At higher K values, one non-admixed group was observed for the Atlantic while further substructures emerged for the Pacific accessions, where three non-admixed groups were found. Population structure analysis also showed a great degree of admixture between the Atlantic and Pacific populations, and SNPs of the Pacific non-admixed genetic groups were mostly introgressed into the Atlantic individuals but the contrary was rarely observed. The results of principal component analysis and Neighbor-Joining Hierarchical Clustering were consistent with the results from Structure and provided a measure of genetic relationships among individual genotypes. The Pacific group has a lower genetic diversity and a higher rate of inbreeding than the Atlantic group. These results suggest that the Pacific coconuts of Colombia belong to the pre-Columbian population found on the Pacific coast of Panama and Peru. If it had been introduced after Columbus (as in Mexico), genetic diversity would have been higher than on the Atlantic coast.

## Introduction

The coconut palm (*Cocos nucifera* L.; Arecaceae) is one of the most economically important palms worldwide due to the versatility of its products, which have been used by humanity since ancient times as a source of food, fuel, medicine, and construction material (Aljohi et al., 2016; Food and Agriculture Organization of the United Nations, 2015; Prades, et al., 2012). Coconut is cultivated extensively in all the tropical countries of the world and is considered one of the most important plant species to guarantee the survival of humankind (Gunn et al., 2011). Many studies have been conducted to characterize the genetic diversity of coconut collections and to understand coconut cultivation history. However, northern South America is an underrepresented region in these studies.

Coconut is a hermaphrodite, perennial diploid (2n = 32) tree and the sole species of the genus *Cocos*. Draft genomes have been published at the scaffolds level, suggesting a genome size of about 2.1Gb (dwarf coconut) to 2.4 Gb (tall Hainan coconut) and a complex genome structure composed of 50-70% of repetitive sequences, chromosome fractionation and duplication followed by rearrangements (Xiao et al., 2017; Yang et al., 2018; Latican et al., 2019). Archeological evidence dates wild coconut fossils to ca. 60 MYA in the Americas (Gomez-Navarro et al., 2009), however coconut origin is still largely controversial. Today the prevalent hypothesis considers *Cocos nucifera* has American ancestors, but its lineage probably became extinct on the continent until it was introduced during the late Holocene, but before Columbus (Baudouin et al., 2014).

Coconut selection by cultivation, hybridization and introgression in different tropical countries has led to the evolution, dissemination and classification of two well-recognized palm genotypes (Gunn et al., 2011; Geethanjali et al., 2018) brought under cultivation in two separate locations globally, the Pacific Basin and the Indo-Atlantic Ocean. They are identified by a combination of their geographical location, type of reproduction and growth habits. There is a record of prehistoric trade routes and of the colonization of the Americas that offers a rich account of coconut dispersal by humans (Gunn et al., 2011). Gunn et al. (2011) indicated that Indo-Atlantic coconuts were already under cultivation in the southern India subcontinent around 2000 - 3000 years ago base on archaeobotanical (Wheeler et al., 1946), linguistics (Fuller, 2007) and written evidence (Menon and Pandalai, 1958). Also, a coconut fruit fossil with intermediate morphological characters between domesticated and cultivated coconuts was dated with C14 at 1400 years (Lepofsky et al., 1992). At present, this tropical tree crop is distributed across 93 tropical countries (Tang et al., 2006), including Central and South America, East and West Africa, Southeast Asia, and the Pacific Islands, and is grown over 12 million hectares of land (Batugal, 2005).

Worldwide coconut diversity studies have been carried out using RFLP (Lebrun et al., 2005; Meerow et al., 2012), microsatellite (Gunn et al. 2011; Lebrun et al., 2005; Meerow et al., 2012; Geethanjali et al., 2018), and AFLP (Perera et al., 1998; Lebrun et al., 2005 Meerow et al., 2012). All these studies detected high levels of genetic differentiation between Pacific and Indo-Atlantic samples. The Pacific group includes four admixed sub-groups with blurred boundaries: Southeast Asia, Melanesia, Micronesia, Polynesia; and a clearly distinct Panama and Peru group. Dwarf coconut palms also belong to the Pacific and form a genetically uniform group, irrespective of their geographical origin (Batugal et al., 2009). The Indo-Atlantic group includes cultivated coconut palms from Africa and the Atlantic coast of Latin America. They originated from the Indian subcontinent and have extensively bred with coconuts from Southeast Asia, resulting from Austronesian migrations to Madagascar (Lebrun et al., 2005). The clear genetic differentiation between these lineages suggest that coconuts have a long-standing evolutionary presence in both oceans and were brought into cultivation independently in each of these regions. This finding has several important implications for coconut domestication, including two geographical origins of coconut cultivation.

Coconuts lack clear domestication traits making it difficult to trace the origins of cultivation strictly by morphology. Two distinctively different forms of the coconut fruit, known as “Nie kafa” and “Nie vai” -- Samoan names for traditional Polynesian varieties-- are known. The Nie kafa form is triangular and oblong with a large fibrous husk. The Nie vai form is rounded and contains abundant sweet coconut “water” when unripe. Also, there are “tall” type palms that have evolved naturally and were disseminated by the ocean currents. Tall palms have an unknown origin and are generally cross-pollinated, having long stems and late bearing fruit, and can be of the “Nie kafa or Nie vai type” (Harries 1981; Gunn et al., 2011). The “dwarf” type palms are recognized for being self-pollinated, evidencing important domestication features such as short stems, high productivity in terms of nut production, and low genetic variability. They grow near human habitats and also have inconclusive origin (Latican et al., 2019). Both of these types of palms gave rise to a vast number of coconut populations of pantropical distribution that are poorly characterized at the genomic level, and that are generally identified based on variable morphological and agronomic traits, which are not apparent in juvenile phases (Ribeiro et al., 2013; Guevara & Jáuregui, 2012; Teulat et al., 2000).

High-throughput sequencing (HTS) has enabled the discovery of SNPs throughout the genome, greatly increasing power for detecting neutral and adaptive patterns of variation (Taranto et al., 2016; Torkamaneh et al 2016). Genotyping by sequencing (GBS) is considered an efficient and economical method to quickly discover SNPs among several individuals simultaneously (Pavan et al., 2017; Taranto et al., 2016) allowing plant breeders to use these resources for crop improvement. Such an approach has benefited many crops, like watermelon (Nimmakayala et al., 2014), cowpea (Xiong et al., 2016), rice (Xu et al., 2012), and spinach (Shi et al., 2017) among many others.

Despite the central importance of coconut as a widely distributed and cultivated palm around the world little is known about the genomic diversity of this species. Here we present a comprehensive catalogue of approximately 27000 single nucleotide polymorphisms (SNPs) in coconut, based on Genotyping by Sequencing (GBS) of a collection of 125 individuals representing wild and commercial varieties that are cultivated around the globe and their hybrids. We aim to determine the genetic identity and diversity of cultivated coconut in northern South America. Our work informs management practices for coconut germplasm resources, provides a better understanding of the genetic diversity and population structure of coconut populations, and adds to the understanding of how coconut palms evolved under domestication.

## Results

### Genotyping of coconut accessions

We performed Genotyping by Sequencing (GBS) analysis of at least four varieties of wild and commercially important and widely cultivated coconut. A total of 125 coconut individuals were obtained from main coconut producing departments in the Atlantic and Pacific Colombian coasts. Three of the non-admixed cultivars are from the Pacific coast, and one from the Atlantic coast. We obtained approx. 367 millions of raw reads for all individuals analyzed in this study (average 3.2, standard deviation 1.2 millions of raw reads per individual; Tables S1, S2; NCBI Bioproject PRJNA579494; NCBI SRA Accessions SRR10345275-SRR10345401). FastQC results indicated that reads were of high quality across the entire 150bp length. Because GBS datasets often have high levels of missing data, filtering parameters can have a major impact on the overall size of the dataset. After filtering loci, the dataset resulted in 125 individuals and 27600 SNPs markers that had a minor allele frequency > 0.01% (Fig. S2), mean missing data 8.57% and standard deviation 3.5%. After pruning for linkage disequilibrium (LD), 13900 unlinked SNPs of 27600 with *r*^2^ < 0.3 were kept for further analysis. Eighteen recent hybrids were detected, all belonging to the Atlantic coast populations (Fig. 1).

**Figure 1.**
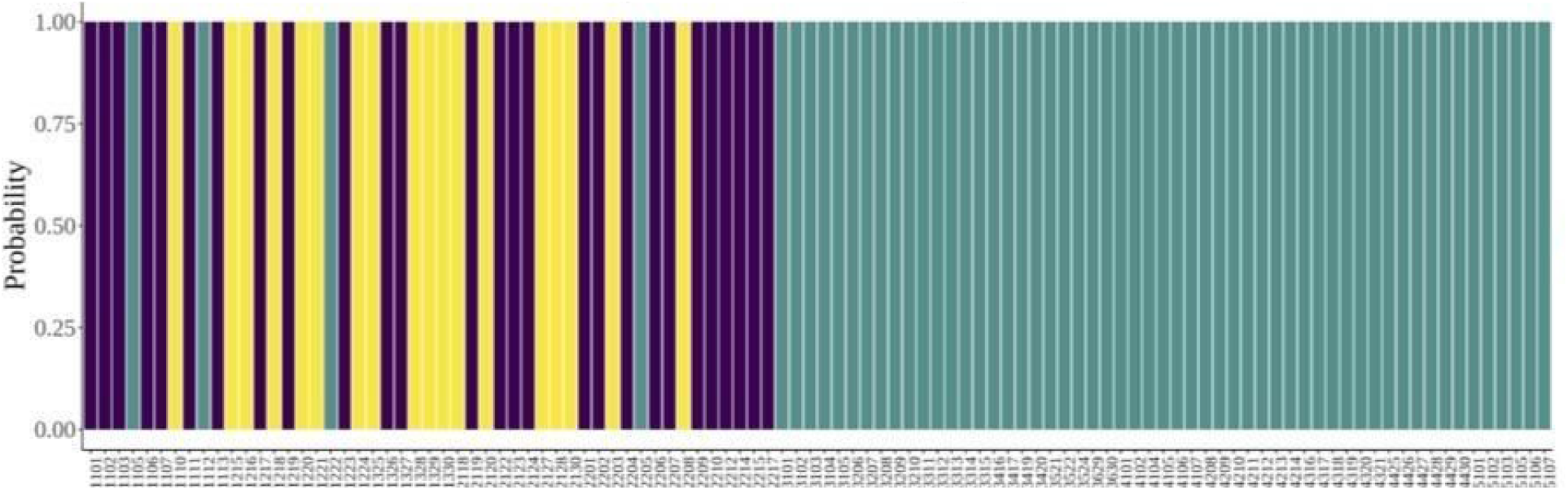
Recent hybrids detection using snapclust function in adegenet (Jombart, 2008). Recent hybrids detection using snapclust function in adegenet (Jombart, 2008). A vertical bar represents each accession. Each color represents either a pure parental (green or purple) or a hybrid (yellow), and the length of each colored segment in each vertical bar represents the proportion contributed by the parents.

### Genetic diversity and differentiation of Northern South America coconut

First we used the larger dataset (125 accessions and 27600 SNPs markers) to calculate summary population statistics for each sampling site (Antioquia, Córdoba, Nariño, Cauca, Chocó) and coastal regions (Atlantic and Pacific). FST calculations between sampling sites and coastal regions, observed heterozygosity, expected heterozygosity and inbreeding coefficient averaged across all loci were calculated. Expected and observed heterozygosity (He, Ho) and inbreeding coefficients (FIS) within Pacific and Atlantic regions are comparable. Mean observed heterozygosity was 0.1654 for the whole collection, 0.0678 for the Pacific and 0.2630 Atlantic dataset. Average He, Ho per coastal region were lower for the Pacific than the Atlantic; while FIS was higher in the Pacific (Fig. 2, Fig.S3).

**Figure 2.**
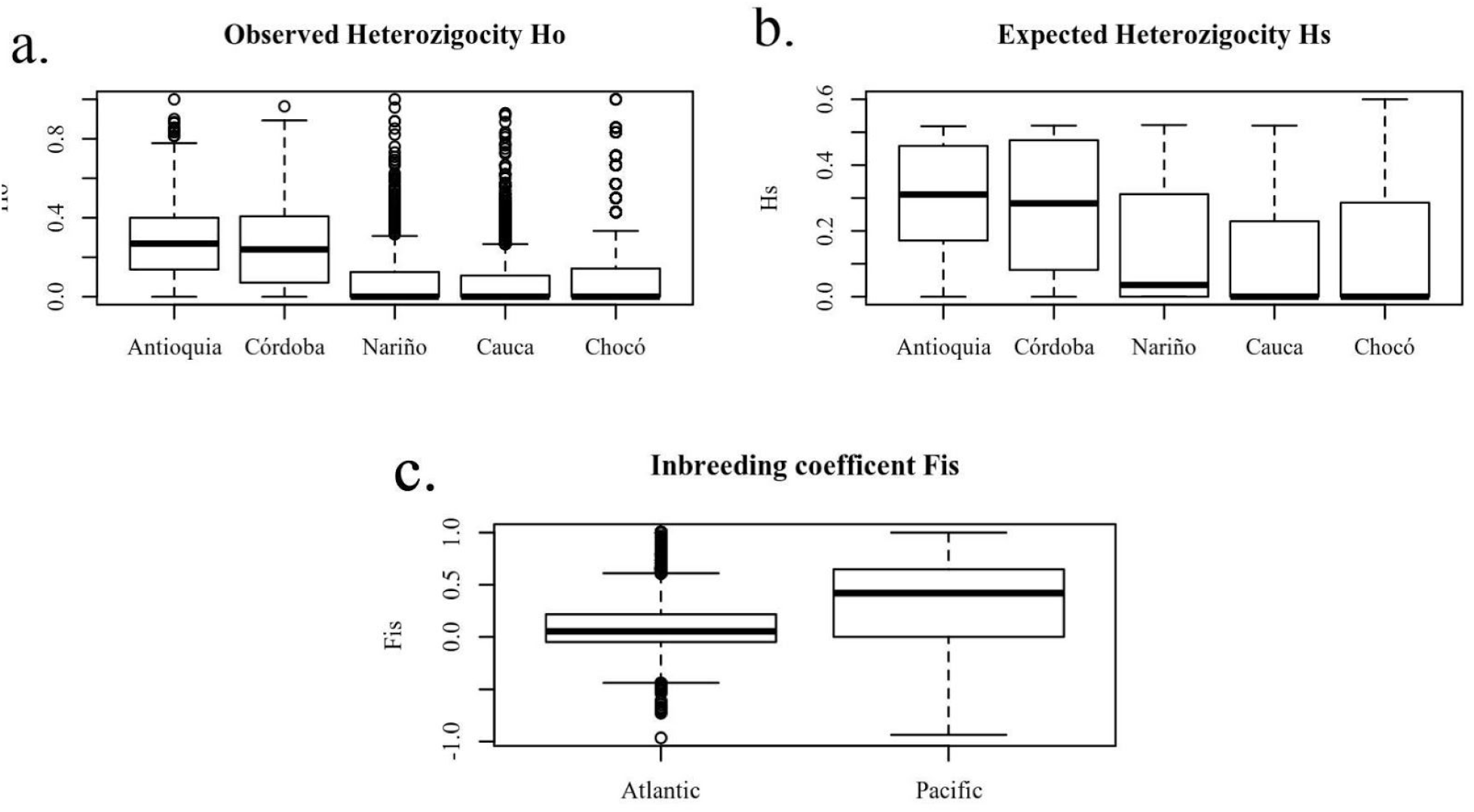
Population statistics per loci calculated using the “*basic.stats*” function of Hierfstat 0.04-30 R package and grouped by coastal region and sampled site. a. Observed heterozygosity per sample site. b. Expected heterozygosity per sample sites. c. Inbreeding coefficient per region. Boxes represent interquartile range, bars represent maximum and minimum values and the points represent potential outliers.

A pattern of genetic differentiation between the two coasts Atlantic and Pacific was found. Also, FST within samples from the same coast are smaller than FST of samples from different coasts, for example, Nariño and Cauca FST (0.0153 - 0.0167), and Nariño and Chocó FST (0.0821 - 0.0844). FST do not overlap indicating differentiation within the Pacific that are an order of magnitude smaller than the FST of Nariño with Antioquia (0.4001 - 0.4073) and Nariño with Córdoba (0.3519 - 0.3578). Lower levels of population divergence were detected in the full data set for comparisons within Atlantic and Pacific populations. The highest genetic divergence was found between Antioquia and Nariño’s populations (FST = 0.400-0.407). Meanwhile the closest genetic divergence was found between Nariño and Cauca’s populations (FST = 0.0153-0.0167) (Table 1). Global FST suggest larger genetic divergence between Pacific and Atlantic coasts (0.3827 p<0.001) rather than the five-group comparison (0.2925, p<0.001).

**Table 1.** Pairwise Loci FST per sampled site in Northern south America Coconut populations. FST was calculated using Nei’s pairwise FST with Hierfstat in R. Pairwise comparisons for each loci between sampled sites were calculated, then a confidence interval [2.5% – 97.5%] was computed using 1000 bootstraps.

|  | Córdoba | Nariño | Cauca | Chocó |
| --- | --- | --- | --- | --- |
| Antioquia | 0.0312-0.0321 | 0.4001-0.4073 | 0.3808-0.3869 | 0.3321-0.3384 |
| Córdoba |  | 0.3519-0.3578 | 0.3495-0.3544 | 0.2877-0.2931 |
| Nariño |  |  | 0.0153-0.0167 | 0.0821-0.0844 |
| Cauca |  |  |  | 0.1204-0.1228 |

We detected FIS ranges from −0.5 to values close to 1 (Fig. 2, Fig. 3S). Atlantic coconut populations are predominantly exogamous due to cross-pollination and those of the Pacific are predominantly inbred due to self-pollination. Global heterozygosity is less than 0.2, which partly reflects the inbreeding habits of most of the Pacific palms and some of the Atlantic (Fig. 2, S3c). Inbreeding coefficients for Antioquia and Cordoba palms had similar positive values close to zero. Pacific populations (Cauca, Nariño and Chocó) had inbreeding coefficients close to 0.5 (Fig. 2c, S3c). Atlantic populations had higher heterozygosity than those of the Pacific (Fig. 2a-b, S3a-b).

**Figure 3.**
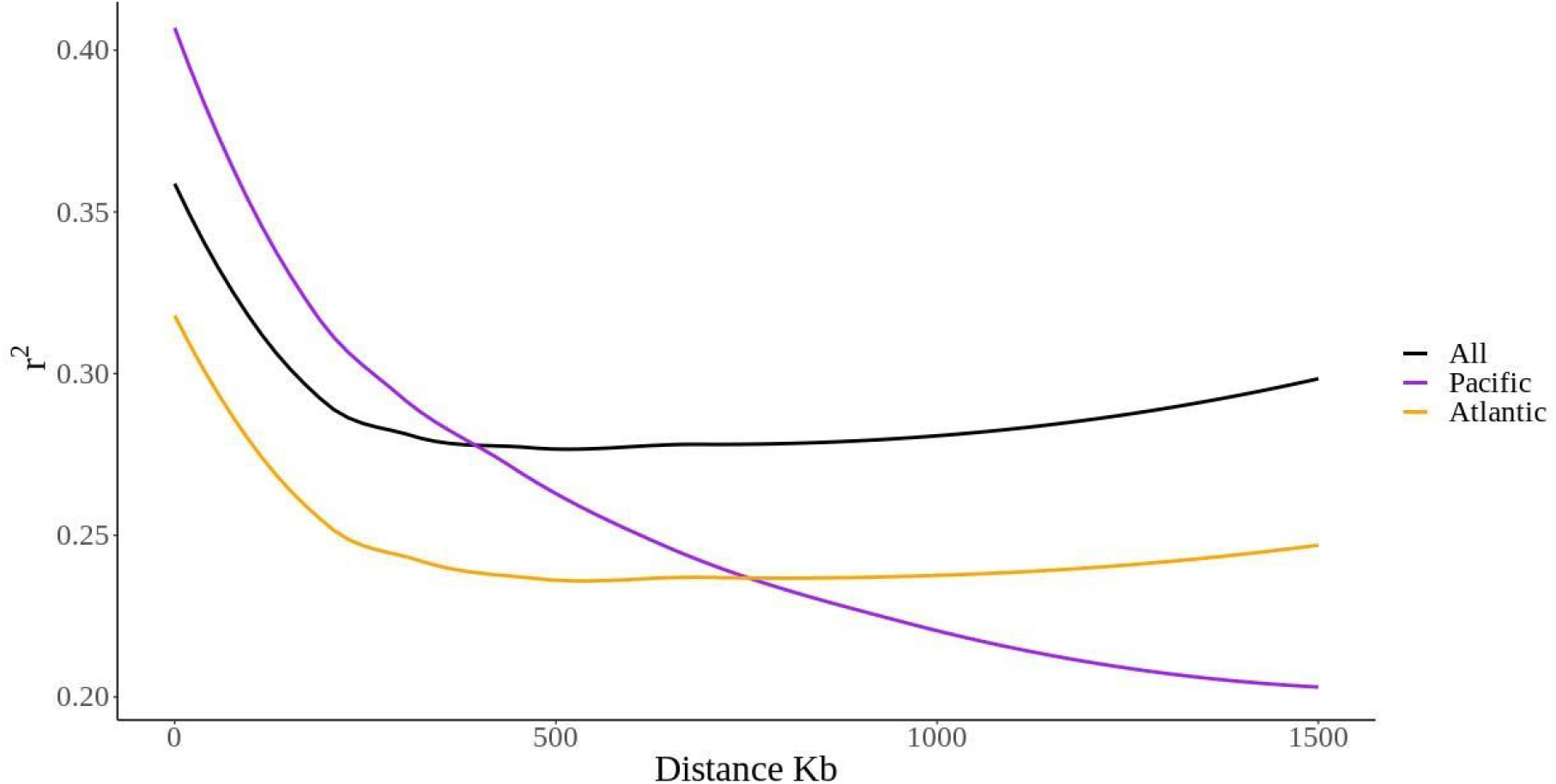
Linkage disequilibrium (LD) graph of coconut grown in Northern South America. LD is determined by squared correlations of allele frequencies (r2 < 0.3) against distance between polymorphic sites, color-coded as follows: all accessions (black), accessions from the Pacific coast (purple) and accessions from the Atlantic coast (orange).

### Linkage disequilibrium

LD was estimated between all SNP markers over the 125 coconut accessions (https://doi.org/10.6084/m9.figshare.13020635). To assess the extent of LD decay, the estimate of r2 for all pairs of SNP loci linked on the same scaffold in the reference genome (Hainan Tall), was calculated. The LD decay was evaluated for the whole dataset and for the accessions from the Atlantic and the Pacific separately. The LD decay observed for the pacific was relatively slower in 1500Kb than in the Atlantic. The squared correlations of allele frequencies r2 of the Pacific stabilizes approximately at 1500 Kb, while for the Atlantic we do reach stabilization approximately at 250Kb (Fig 3).

### Population structure of Northern South America cultivated coconut

To understand population structure of coconut cultivars in northern South America we used several approaches, Discriminant Analysis of Principal Components (DAPC), Minimum Spanning Network (MSN) and Bayesian clustering methods (Fig. 4 and 5). Accessions tend to group by either Pacific or Atlantic coasts. The MSN based on Euclidean distances between genotypes, shows the Atlantic accessions form two close subgroups, a first with a mix of accessions from Antioquia and Córdoba, and a second one with a mix of accessions of Antioquia, Cordoba and some of the Nariño accessions (Fig. 4a). Three groups are also formed in the Pacific coast using the MSN analysis. The first group is closer to the Atlantic accessions, a second one is distant and includes individuals from Cauca, Nariño and Chocó and last, a smaller group is formed with individuals from Nariño and Chocó (Fig. 4a). DAPC was done with the first 39 eigenvalues representing 81.33% of the variation (Fig. 4b). This analysis splits genotypic data into three groups divided according to the two core groups (Pacific and Atlantic), with one single group shown within the Pacific accessions and two distinct groups formed from the Atlantic accessions, corresponding with the populations from Córdoba and Antioquia (Fig. 4b).

**Figure 4.**
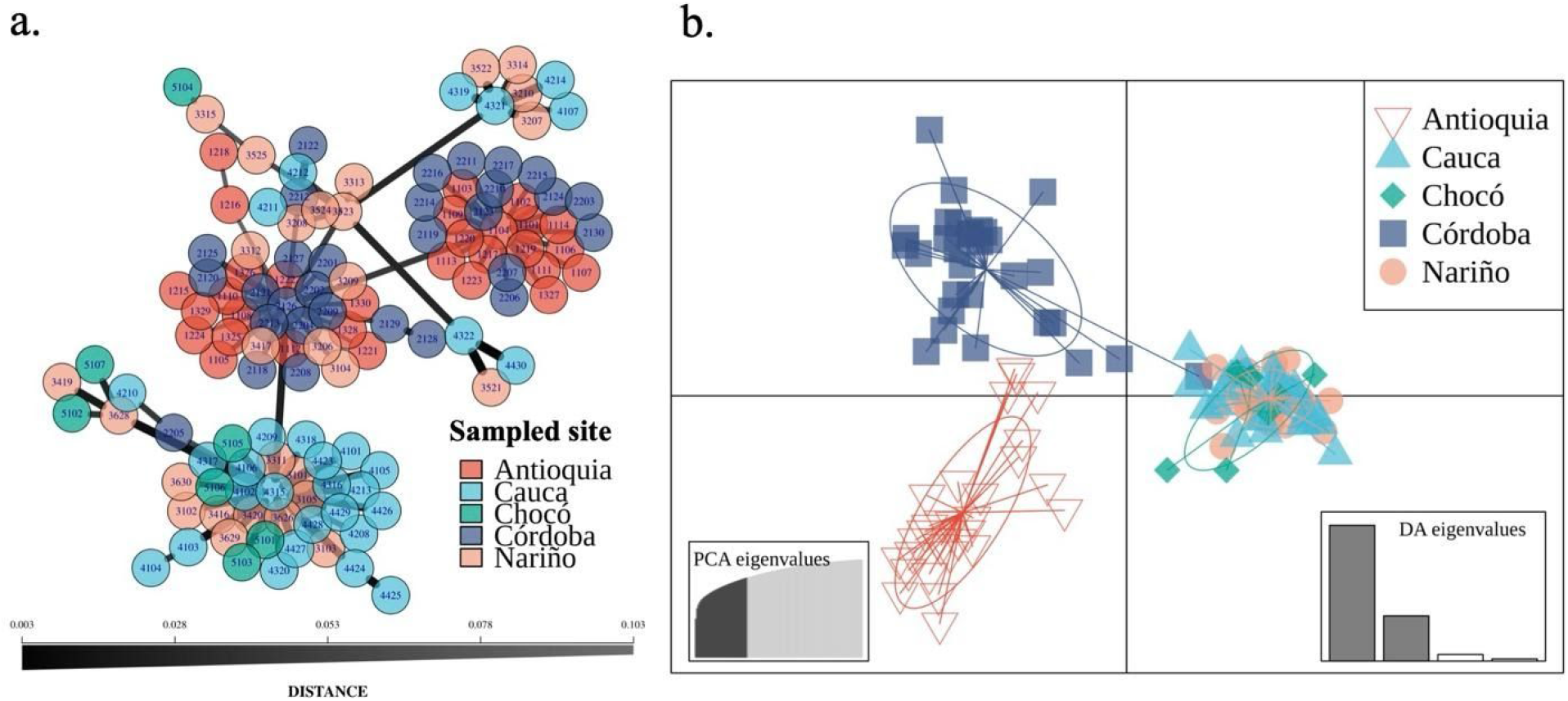
Coconut genetic groups formed in Northern South America using Minimum Spanning Networks (MSN) and Discriminant Principal Component Analysis (DPCA). a. Minimum spanning network based on Euclidean distances, numbers in each circle are the codes for collected individuals (refer to Table S1). b. Discriminant Analysis of Principal Components (DPAC) of 39 principal components explaining 81.33% of the variance and two discriminant axes, ellipses explain 67% of variation.

**Figure 5.**
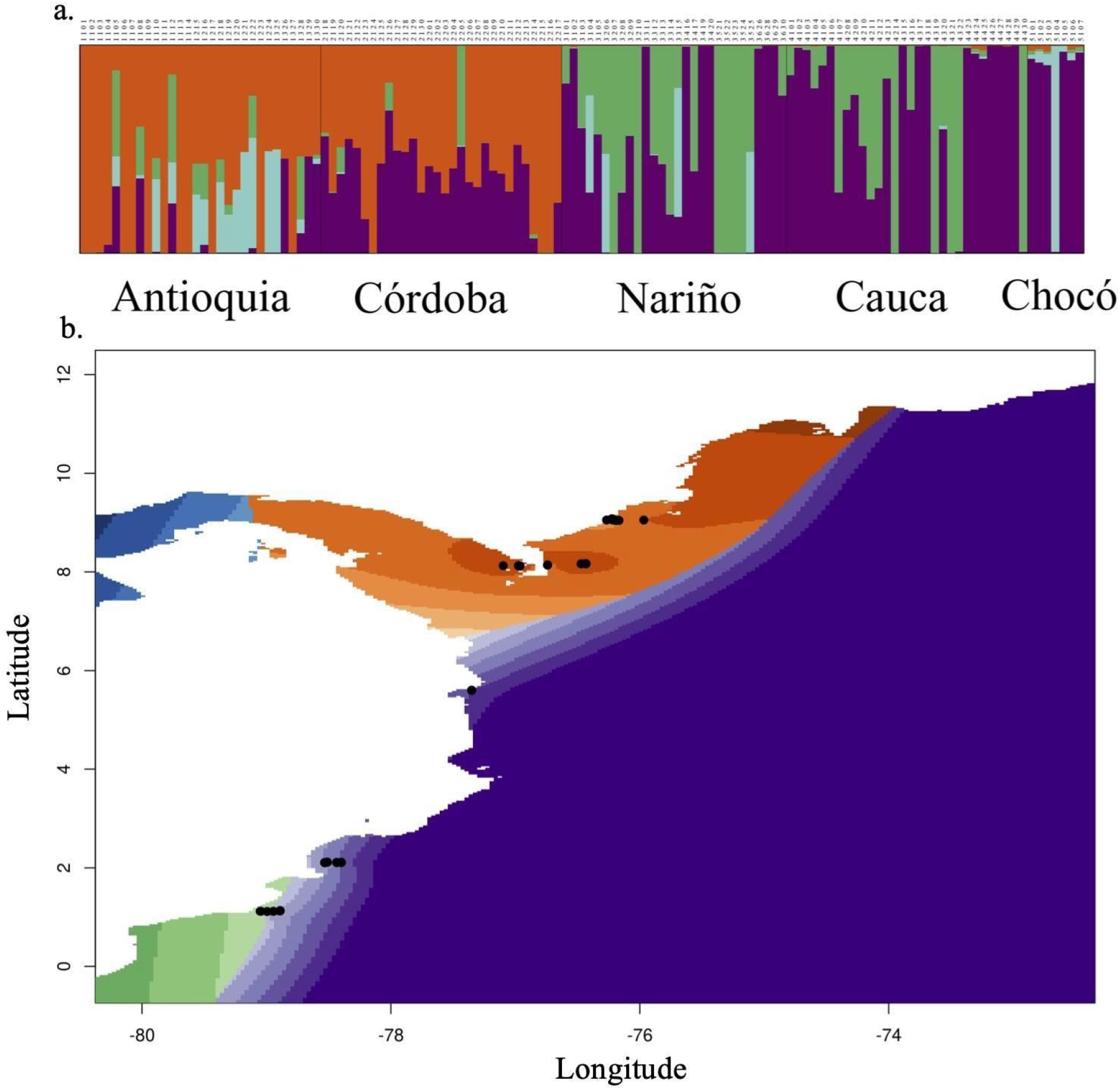
Population structure of northern South America cultivated coconuts using a larger number of SNPs recovered by STACKS and Spatial Interpolation of Ancestry Coefficient Maps. a. Population structure analysis using STRUCTURE including 26700 SNPs and 125 individuals. A vertical bar represents each accession. Each color represents one ancestral population, and the length of each colored segment in each vertical bar represents the proportion contributed by ancestral populations. b. Spatial interpolation of ancestry coefficient maps.

Bayesian clustering algorithm implemented in Structure (Hubisz et al., 2009) was used to estimate ancestry proportions for each coconut accession. Evanno’s test showed that four populations (K = 4) represented the best number of clusters for 125 coconut accessions and 27600 SNPs (Fig. 5; Fig. S4). However, we performed analysis at K = 2 and K = 3 too (Fig. S5). A K = 4 was also confirmed by the MSN in a reduced dataset with missing data rate < 5% (27600 SNPs and 58 of 125 total accessions). Population structure results at this optimal K is also consistent with DAPC analysis (Figs. 5a). All analysis suggests two primary clusters and a subtle substructure within. As shown in Fig 4 and Fig. S5 from K = 2, to K = 4, the Atlantic group was clearly separated from the Pacific group. At K = 4 a large portion of accessions showed genetic admixture, but a group of SNPs are exclusive either from the Atlantic or the Pacific accessions (Fig. 5a). A second Structure analysis was performed filtering SNPs based on Linkage Disequilibrium (LD) with r2>0.3, a total of 13900 were left of the initial 27600 SNPs. Evanno’s test showed that two populations (K = 2) represented the best number of clusters (Fig. S6a).

### Phylogenetic networks

A phylogenetic network revealed accessions from the Atlantic to be widely separated from and the Pacific. Within the Pacific coast cluster, separation between three groups of palms was found (Fig. 6), and much reticulation among Pacific type palms. In the Atlantic some samples from Antioquia form a well-defined group, but samples from Cordoba seem to have much reticulation. This suggests high levels of hybridization or incomplete lineage sorting among cultivars in this area of Colombia.

**Figure 6.**
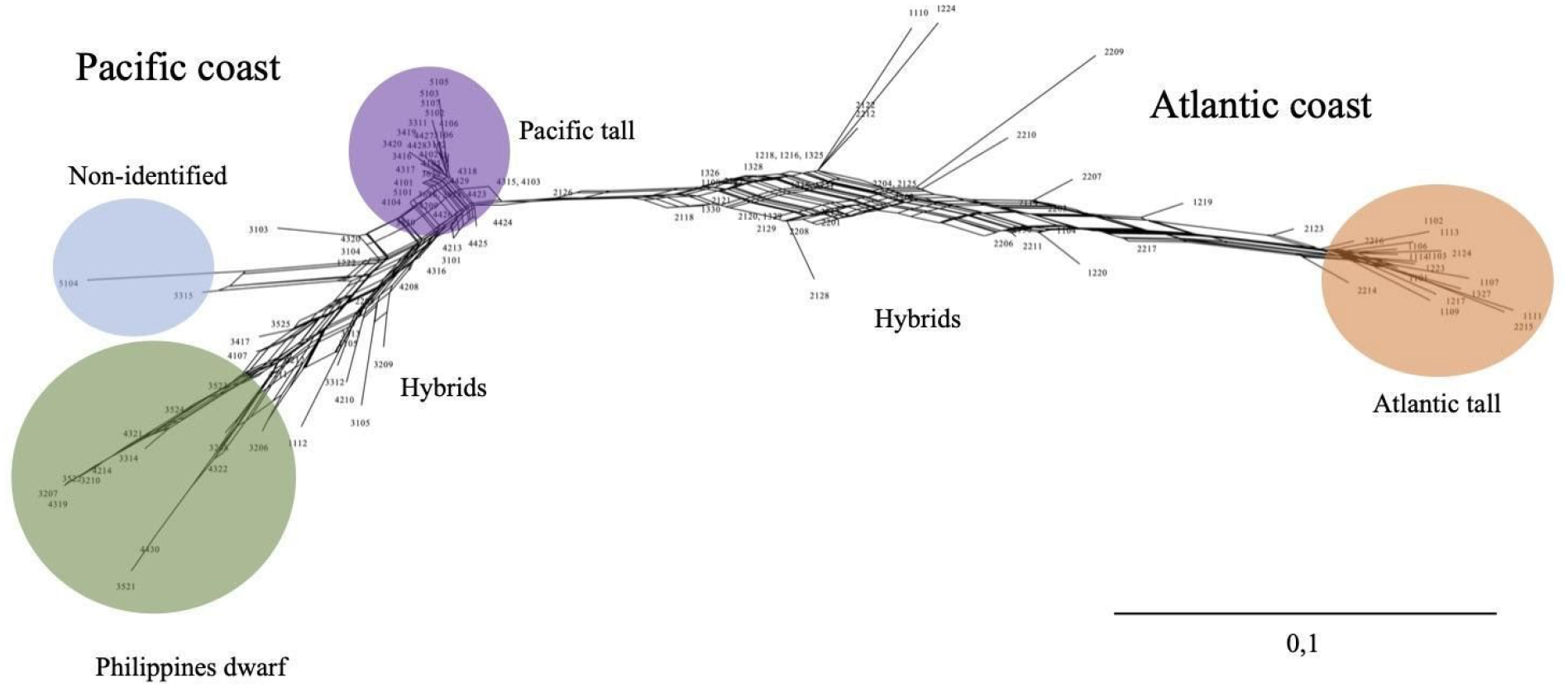
NeighborNet network shows reticulate genetic relationship between accessions. Color coding corresponds to the STRUCTURE clusters of Figure 5. Phylogenetic network calculated using the Neighbor-net algorithm across all individuals. A scale is shown inset. Colors correspond to major accession types shown in Fig. 5.

### Coancestry matrix

Shared ancestry was also analyzed using fineRADstructure, this method derives from utilizing haplotype linkage information focusing on the most recent coalescence (common ancestry) among the sampled individuals to derive a “coancestry matrix” - a summary of nearest neighbor haplotype relationships in the dataset (Malinsky et al., 2018). The analysis shows blocks of related accessions as squares. Darker colors indicate a higher level of coancestry between individuals and the tree depicted at the top of the plot shows relationships among individuals and posterior probabilities as support values on branches (Fig. 7, S7). A first general analysis including the larger dataset with 27600 SNPs and 125 individuals was performed. Two main clusters were formed separating the Atlantic from the Pacific accessions (Fig. S7). In the Atlantic a smaller nested cluster emerged having higher coancestry levels (Fig. S7). Further structure was also found within the Pacific cluster where two sub-groups, one of which had a smaller nested group of accessions from Cauca and Nariño with higher co-ancestry levels (Fig. S7).

**Figure 7.**
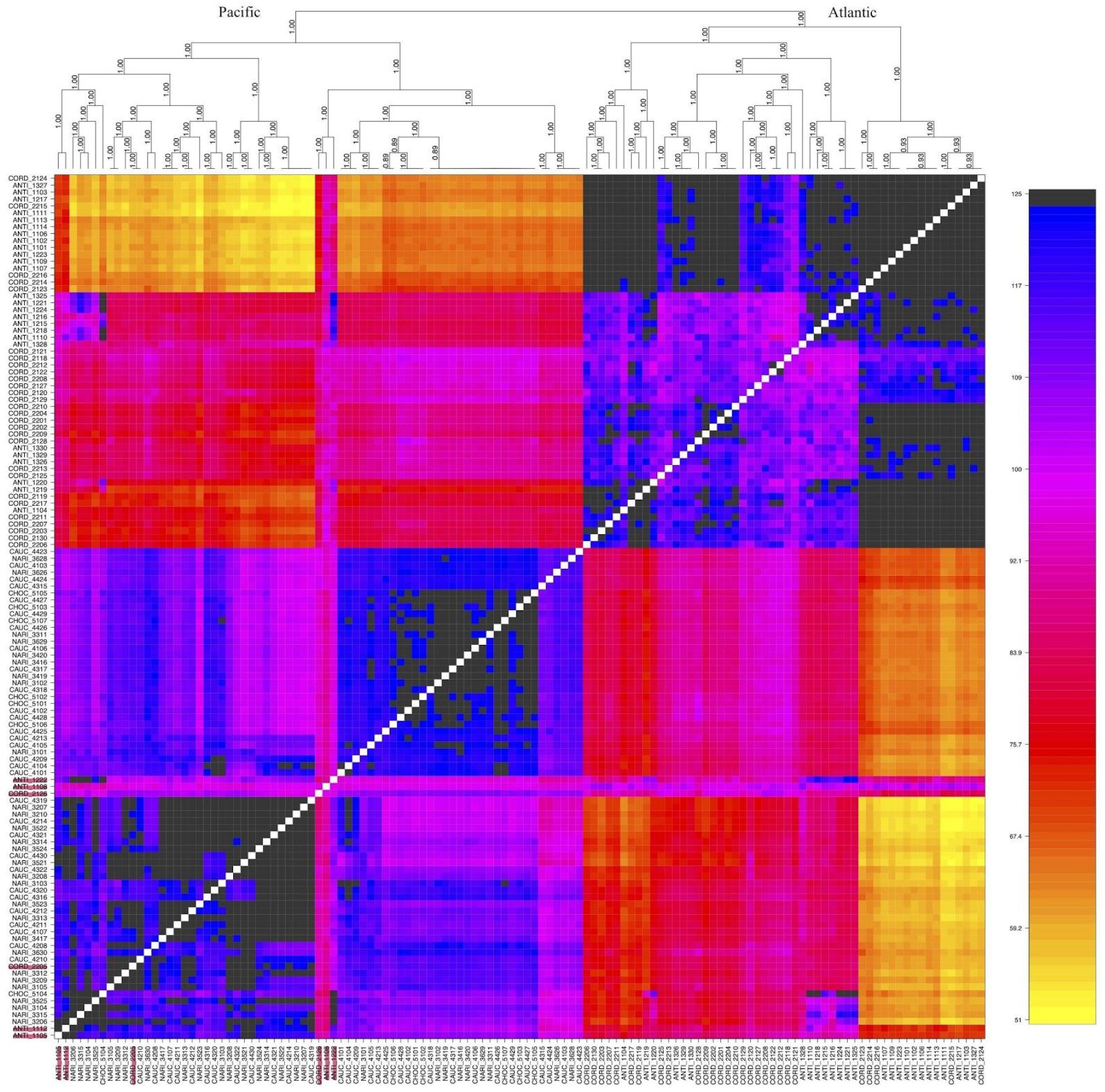
Coancestry matrix for 125 individuals based on 13.900 SNPs after filtering SNPs with linkage disequilibrium (r2 < 0.3) calculated using fineRADstructure. The heat map depicts the high-resolution genetic relationship structure of individuals across *Cocos nucifera* growing in Colombia. Darker colors indicate a higher level of coancestry between individuals. Atlantic and Pacific coast accessions are separated and there is a level of structure amongst 125 individuals. The tree depicted at the top of the plot shows relationships among individuals and posterior probabilities as support values on branches, further information can be found in Malinsky et al. (2018). Highlight accessions in pink and in both axes represent Atlantic accessions.

Using the LD filtered dataset (r2 > 0.3 13900/27600 SNPs) a second analysis was performed. Two main clusters were also recovered in this analysis by coast, all of which have relatively higher coancestry levels and nested substructure, in comparison to the first analysis (Fig. 7). Six nested clusters with the highest coancestry levels are recovered among accessions, including some from the Atlantic and some from the Pacific coasts (grey color in Fig. 7).

### Relationships among morphological data, agronomic traits and inbreeding coefficients

To better understand the phenotypic diversity of the coconut populations found in Colombia, in relation to results obtained with molecular markers, we measured morphological and agronomic variables in all accessions during 2017-2019. A Principal Component Analysis (PCA) was performed. Phenotypic data was analyzed together with the inbreeding coefficient (FIS) calculated for each individual using Vcftools (Danecek et al. 2011).

PCA included mean values from five morphological characters from palms (rachis length, plant height, number of leaflets per plant, stem circumference and the ratio between the equatorial and polar diameter for the nuts). The first and second principal components account for 35% and 21% of the total variation, respectively (Fig. 8a). We found a clear differentiation between the Pacific and Atlantic groups, explained in the Pacific mainly by the FIS coefficient, while in the Atlantic most morphological characters define this group (Fig. 8a). A second PCA analysis was performed using average harvest data from three seasons (2017-2019) and the inbreeding coefficient (FIS). This PCA shows cluster separation between the Atlantic and hybrids with intermedia morphological values. The first and second principal components account for 55 and 15% of the total variation, respectively (Fig. 8b). Pacific accessions formed two separate clusters explained by the FIS and the number of coconuts (N.coconuts), racemes (N.racemes), water (W) and the fiber weight (FW) produced by each palm (Fig. 8).

**Figure 8.**
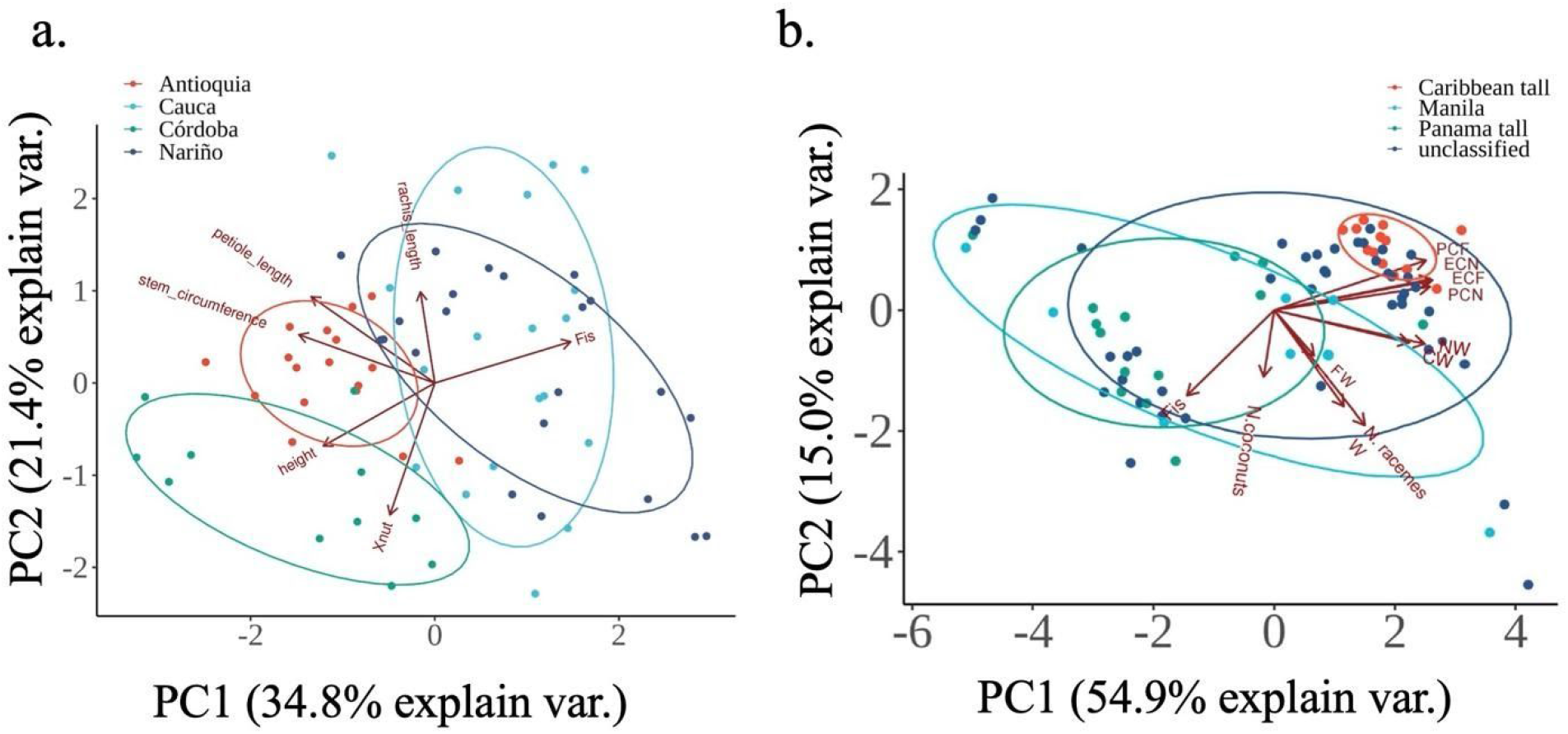
Discriminant Principal Component Analysis of morpho-agronomic, average harvest data and inbreeding coefficients. (a) PCA of selected morpho-agronomic data (rachis length, plant height, number of leaflets per plant, stem circumference and the ratio between the equatorial and polar diameter for the nuts (Xnut) and inbreeding coefficient). (b) Average harvest data from several seasons (2017-2019). Polar circumference of the fruit (PCF), equatorial circumference of the fruit (ECF), polar circumference of the nut (PCN), equatorial circumference of the nut (ECN), water volume (W), copra weight, (CW) and fiber weight (FW) and Inbreeding coefficient (FIS).

## Discussion

Reduced representation libraries from high-throughput sequencing (HTS) allow analysis of SNPs throughout the *Cocus nucifera* genome, providing both neutral and putatively selected markers. Genotypes were successfully obtained for 125 samples from five different geographical regions in northern South America. This is the first genome wide approach used in coconut to discover SNPs, filling an important gap in the published genomes (Xiao et al., 2017; Latican et al., 2019). In this study, we provide a large genome variation data set for cultivated coconut grown in Colombian coastal regions that have not been previously characterized at the molecular level. Population structure not only supports the hypothesis that there is a strong genetic break between Atlantic and Pacific populations in northern South American but also provides finer structures within the Pacific coast. This pattern corroborates previous hypotheses for worldwide genetic diversity of coconut (Gunn et al., 2011).

We recovered a total of 27600 SNPs along the genomes of 125 accessions of coconut. Sequence variations between ‘Hainan Tall’ and ‘Catigan Green Dwarf’ (CATD) genomes included 57872 SNPs in intergenic regions, 21066 in genic regions and 5552 in exonic regions. This study recovered 32.69% of the total number found in Xiao et al. (2017) and Latican et al. (2019).

All approaches used reveal there is pronounced genetic structure in coconut accessions from Colombia even though both coasts are geographically close to each other. The structure analyses, principal component analysis, discriminant principal component analysis and minimum spanning networks are all consistent with 4 groups explaining the ancestry of these populations. FST analyses suggest the differentiation between Pacific and Atlantic coasts is more important in explaining the variance structure, yet differences between these components of ancestry within regions are significant. Here, the sampling strategy was to select individuals trying to cover all different cultivated coconut varieties present in each zone due to absence of information about specific cultivars and varieties that are grown in each zone. We found a pattern of genetic differentiation between the two coasts Atlantic and Pacific. This pattern was initially revealed at a global scale (Gunn et al., 2011) and is now confirmed at a local scale using genomics in northern South America. Pacific coconuts are showing genetic substructure and higher inbreeding coefficients (Fig. 2–7). This level of differentiation suggests long-term evolutionary divergence between two subpopulations, with independent origins of naturally occurring and cultivated coconuts from within each lineage (Table 1).

Our Structure (K = 4) data suggest extensively admixed genotypes for the Atlantic coast of Colombia. However, we identified four non-admixed genotypes including most of the total genetic variation in the region (Figs. 3, 4). A group of SNPs are exclusive to the Atlantic accessions while SNPs from the Pacific accessions were found in admixed accessions from the Atlantic (Figs. 3–5), indicating gene flow could be exclusively unidirectional or very limited from the Atlantic to the Pacific populations. We could be overestimating K, due to the widespread pattern of admixture; plots with K=2 and K=3 are also included in supplemental material (Fig. S5).

Here, non-admixed Atlantic tall coconuts showed phenotypic similarity (Fig. 8) and grouped together in coancestry analysis (Fig. 7) representing what we consider to be the wild type coconut genotype from the Atlantic. Sixteen non-admixed accessions have SNPs exclusive to the Atlantic Tall type and very few of these SNPs were found in accessions from the Pacific (Fig. 5). They display characteristics of the Atlantic wild type described by Harries (1981). Historical records suggest that the introduction of coconut palms to the Atlantic coasts of America originated from the Cape Verde Islands, where they had been previously introduced from Mozambique by Portuguese sailors after 1499. Introductions were also made to the coasts of West Africa during the 16th century (Harries, 1992; Schuiling and Harries, 1994). However, coconut fossils in the Atlantic coast of Colombia have been dated to 60 MYA (Goméz-Navarro et al., 2009).

In this study, Dwarfs coconuts were identified because most of them have a high inbreeding coefficient (> 0.9), short internodes (< 1m) and the origin of seeds seem to be Southeast Asia according to the information provided by Colombian farmers. We consider these accessions to be of the Philippine type Dwarf, a non-admixed, distinctive coconut cultivated type (Table S1, see in Fig. 5 green color). A recent molecular SSR study has shown differences in allele frequency between Dwarf and Tall accessions, as in this study. It has been suggested Dwarf types originated from Tall through domestication that likely occurred in Southeast Asia. Domestication took place through autogamy followed by a series of allele fixations in a random subset of ancestral Talls (Perera et al. 2016; Latican et al., 2019). Dwarf coconuts show the lowest levels of variation in our samples. According to Perera et al. (2016) who also found reduced diversity in Dwarfs, an indication of a population bottleneck. They also suggested that this is a consequence of a series of mutations, resulting in the female flowering phase occurring earlier and lasting longer. As a result, in Dwarfs, female and male flowering coincide, strongly aiding self-pollination.

Admixed accessions from the Philippe type share many SNPs with a second group we believe to be the “Pacific Tall” also known as the “Panama Tall” type, reported from Panama, Ecuador and Peru and now also confirmed for Colombia (Bauding and Lebrun, 2009). The Panama Tall is proposed to be descended from cultivated populations because of its low genetic diversity, possibly a product of a genetic bottleneck; however, the low genetic diversity could also be explained by a natural founder event. The Pacific Tall has been hypothesized to descend from an ancestral Southeast Asian population (Bauding and Lebrun, 2009; Baudouin et al., 2014) (in Fig. 5 purple color). They suggest it must have been brought to the Americas, because the distance from the Philippines to Panama prevents unaided drifting. Our findings support a wide distribution of the Pacific Tall along the pacific coast of Panama and northern south America that predates Columbus, casting shadow to Baudouin et al. (2014) hypothesis. A recent publication has also shown gene flow of Native American human populations predated the pacific island populations settlement (Loannidis et al., 2020) bringing back the debate about the possible American origin of coconut.

Most Tall coconuts from the Colombian Atlantic coast were admixed with one, two or three distinctive group of SNPs from the Pacific, they are Tall coconuts predominantly cross-pollinating usually having a low to negative inbreeding rate FIS: [−1.07, - 0.6], height: [5 - 18.5m]. SNPs belonging to the Pacific Tall type are also widely admixed with accessions from both the Atlantic and the Pacific coast, but non-admixed individuals are only found in the Pacific (FIS: 0.39-0.79, height: 1.5 - 10m). A third group of SNPs of unknown origin was found in a single non-admixed individual collected in Nuquí, Chocó (in Fig. 5 light blue color). Some admixed individuals from both the Atlantic and the Pacific share SNPs with this accession. Looking at the Minimum Spanning Network and the phylogenetic networks this accession seems to be a different type of Dwarf coconut from South East Asia (Figs. 4 and 5). Further studies will have to be performed to find out which accessions represent this group of SNPs that could further clarify the observed genotype.

Besides the admixed accessions found in each coast, in this study we have found a series of recent hybrids (Fig. 1) that were initially used for a first round of analysis and then excluded in a second data round of analysis. Eighteen accessions from the Atlantic were found to be recent hybrids (spontaneous or not). A proportion of these hybrids appeared to be a mix of Dwarfs and Talls with SNPs of the Atlantic Tall type and with the Pacific Tall. They also probably include commercial planting material, which is usually made up of Dwarf × Tall hybrids that have a semi-tall phenotype. Hybrids or Semi-Talls are phenotypically distinct from the two types (Dwarfs and Talls) and can be fixed as with reproductive traits similar to Dwarfs with faster growth (Batugal et al. 2005; Latican et al., 2019). Usually commercial F1 Dwarf × Tall hybrids are produced by emasculating the inflorescences of Dwarf coconuts planted in isolation and applying pollen from selected Talls (Perera et al., 2016).

MSN analysis shows that the Atlantic Talls split in two groups, one including non-admixed Atlantic Talls mostly from Antioquia and a second one containing mostly admixed accessions from Cordoba (Atlantic), grouped with some of the Nariño (Pacific) accessions. To the best of our knowledge this is not shown in any other study. This intermediate group contains admixed accessions with SNPs from the Atlantic, SNPs from Pacific Tall and/or Dwarfs (Fig. 4a). We think Dwarfs and Pacific Talls have been used to produce most of the material that is planted in Cordoba. The MSN also showed accessions from Pacific coast formed an admixed group of mostly Dwarfs and a second formed by non-admixed Pacific Tall accessions (Fig. 4a).

Linkage disequilibrium (LD) has not been mentioned in any other coconut study so far. LD is the non-random association of alleles at different loci and is influenced by various factors. For instance, domestication, population subdivision, and selection can enhance LD in the genome. Generally, LD decays faster in outcrossing species than in self-fertilizing plants (Pavan et al., 2018), and this is what we observed in our data (Fig. 6). Atlantic accessions decay faster at ~250Kb than Pacific at ~1500Kb. The difference in LD decay between the Atlantic and Pacific might be related to the different reproductive systems within *C. nucifera*, and to domestication and selective pressure preserving specific haplotype blocks. LD can be affected by extreme genetic drift in domestication and breeding during evolution. A relatively slow LD decay as the one we observed in inbreed coconut from the Pacific has been observed in other perennial or clonally propagated crops, such as artichoke (Pavan et al., 2018). The rapid LD decay for Atlantic Talls and the high proportion of SNPs in LD (R2 > 0.3, 13900 SNPs) suggest GWAS can be used to inform the breeding of the coconut varieties in Colombia.

We used fineRADstructure to further understand the relationships between the different coconut accessions found in Colombia. After filtering loci based on LD (R2 > 0.3) we manage to recover a strong coancestry matrix with a tree having higher posterior probability values mostly in the branches leading to individuals from the Pacific. In general, this analysis shows the same structure that all other analysis, the accessions group by the oceans where they grow. We were also able to recover a finer and nested structure with this analysis (Fig. 7). In the Atlantic group we find the Atlantic Tall phenotype and a group of what we consider the admixed accessions. In the Pacific we find two clearly distinct groups: the Dwarfs and the Pacific Talls and some nested groups within all having high coancestry values (Fig. 7) but showing same coancestry patterns that SNPs without filtering (Fig. S7).

It has been proposed that coconut was domesticated multiple times (Gunn et al. 2011) and this helps to explain why pacific cultivars are genetically distant from Atlantic ones (Fig. 5). Further work could incorporate more wild individuals from the Pacific coast of Colombia and examine whether cultivated samples from the Pacific Islands present a continuum or a hard genetic discontinuity as might be expected with multiple domestication events.

Lastly, our analysis also included morphological and agronomy data for three different harvest seasons between 2017-2019 and for each sampled site (see raw data here: https://doi.org/10.6084/m9.figshare.13020623). We ran PCA analyses using selected morphology data, average agronomic data and the inbreeding coefficient FIS calculated for each individual. PCA analysis using morphological data shows two groups nearly coinciding with the Atlantic and the Pacific already observed divisions differentiated mostly by FIS and palm height values (Fig. 8a). Using the agronomic data including the average harvest data and the FIS two groups are formed in the PCA analysis. The first component explains the variation in measures related to the fruit shape (Fig. 8b). FIS contributes to the differentiation between Atlantic and Pacific accessions in the first principal component. Also, the Pacific accessions are forming two different subgroups we consider to be the Pacific Tall and the Manila Dwarf along the principal component. These are explained by the number of coconuts in the plant and/or the variation in the mesocarp weight (MW) and the weight of the fruit (FW) (Fig. 8b). Compared to the Atlantic Tall coconuts, the Dwarfs and their hybrids, the Pacific Tall coconuts have smaller fruit (PCF, ECF) and nut size (PCN, ECN) and fiber weight (FW). However, they have a higher amount of liquid endosperm in the fruit (W) (Table S2; https://doi.org/10.6084/m9.figshare.13020623).

## Conclusions

Genotyping by sequencing data has proved useful and reliable for the identification of high-quality SNPs in coconut. We investigated genetic diversity and structure of coconut populations of northern South America. The pattern of variation found by genomic wide analysis indicates the existence of four groups of coconut populations in Northern South America: (1) the Pacific groups with a slow LD decay that includes one or two groups of Dwarf coconuts with higher inbreeding coefficients and the Pacific Talls with high inbreeding coefficients, (2) the Atlantic Talls with high genetic diversity, and fast LD decay phenotypic similarity to the wild type coconut populations. The combination of Bayesian and Hierarchical clustering tools proved to be effective in elucidating population genetic structure of coconut genotypes since the two methods corroborate each other well. Information generated in this study will contribute to the knowledge of coconut genotypes and genetic diversity present in Colombia. Red ring disease in coconut and other cultivated and native palms in the Neotropics have reached epidemic levels. Knowledge of the northern South America genetic structure of the coconut, including regions with augmented levels of genetic diversity, may ultimately prove useful in targeting source populations for disease resistance and other crop improvement traits. This work is a first step towards future genome-wide association mapping studies and the identification of SNP markers able to enhance the precision breeding for horticultural traits in cultivated coconut palms.

## Methods

### Plant materials and experimental design

A total of 127 coconut mature individuals were selected from the main coconut producing departments on the Atlantic and Pacific coasts of Colombia. Two sampled sites in the Atlantic coast included: 1) 30 accessions from Córdoba, municipalities of Puerto Escondido (locations: Mucuna, El Paraiso, La Union and Puerto Alegre) and Moñitos (locations: Pueblito, La Rada, Behiacoita and El Destino). 2) 30 accessions from Antioquia, municipalities of San Juan de Urabá (locations: Uveros y La Balsilla) and Arboletes (locations: La Fortaleza y El Destino). Three sites were located in the Pacific coast: 1) 30 accessions from Nariño, municipality of Tumaco (locations: San Jose del Guayabo, Tablón Dulce, Chagui, Buenos Aires, Rosario and Gualayo). 2) 30 accessions from Cauca, municipality of Guapi (locations: Playa Blanca, Quiroga, Preba, Obregón) and 3) 7 accessions from Chocó, Municipality of Nuquí (location: Coquí). Selection of accessions was made trying to cover all different cultivated coconut varieties present in each zone. There was no official available information about specific cultivars and varieties that were grown in each zone. 116 individuals were geo-referenced and marked in the field (Table S1).

The Minimum List of Coconut Descriptors (IBPGR, 1992) was used to select a suite of qualitative and quantitative variables that could be used to measure morpho-agronomic characters in coconuts. Analyzed variables included: palm shape, trunk circumference (cm), plant height (m), petiole length (cm), rachis length (cm), number of leaflets, fruit shape, nut shape, number of coconuts per palm, fruit polar circumference (cm), fruit equatorial circumference (cm), nut polar circumference (cm), nut equatorial circumference (cm), fruit weight (g), nut weight (g), copra weight (g), fiber weight (g), and volume of coconut water (ml) (Table S2 full datasets are available in Figshare: https://doi.org/10.6084/m9.figshare.13020623). Sampled coconuts were visualized through a Principal Component Analysis (PCA) in R.

### DNA extraction, library preparation and sequencing

DNA extraction was performed from fresh leaf tissue collected in the field. A combination of CTAB (Hexadecyl Trimethyl Ammonium Bromide) extraction method (Murray & Thompson, 1980) with Epoch Life Science® (Epoch Life Science Inc. Missouri, Texas, USA) purification columns were used to extract and purify DNA. DNA quality was determined by spectrophotometry using Nanodrop 2000 (Thermo Fisher Scientific®, Waltham, Massachusetts, USA) and DNA concentration through fluorometry using Qubit 3.0 (Life Technologies® Carlsbad, California, USA).

Library construction and sequencing was outsourced with LGC Genomics (Berlin, Germany). Initially a pilot experiment using 12 samples (3 accessions x 4 populations/2 Pacific coast and 2 Atlantic coast) was performed to find out which set of restriction enzymes work better to recover more Single Nucleotide Polymorphisms (SNPs) in coconut. Between 100 and 200 ng of genomic DNA was digested using 2 units of the MslI restriction enzyme (New England Biolabs ®, Ipswich, Massachusetts, USA.) and using 1 unit of NEB4 buffer in 20 μl of volume and for 2 hours at 37 °C. The restriction enzyme was inactivated by incubation at 80 °C for 20 min. The same procedure was used for double digestion with the PstI-MspI enzymes (New England Biolabs ®, Ipswich, Massachusetts, USA).

For the ligation reaction, 10 μl of each restriction digest were transferred to a new 96 well PCR plate, mixed on ice first with 1.5 μl of one of 96 inline-barcoded forward blunt Adaptors (pre-hybridized, concentration 5 pmol/μl), followed by addition of 20 μl Ligation master mix (containing: 15 μl NEB Quick ligation buffer, 0.4 μl NEB Quick Ligase, 7.5 pM pre-hybridized common reverse blunt adaptor). Ligation reactions were incubated for 1h at room temperature, followed by heat inactivation for 10 min at 65 °C. For library purification, all reactions were diluted with 30 μl TE 10/50 (10mM Tris/HCl, 50mM EDTA, pH: 8.0) and mixed with 50 μl Agencourt XP beads, incubated for 10 min at room temperature and placed for 5 min on a magnet to collect the beads. The supernatant was discarded, and the beads were washed two times with 200 μl 80% ethanol. Beads were air dried for 10 min and libraries were eluted in 20 μl Tris Buffer (5 mM Tris/HCl pH: 9.0).

For library amplification, 10 μl of each of the 96 Libraries were separately amplified in 20 μl PCR reactions using MyTaq™ (Meridian Bioline, Memphis, Tennessee, USA) and standard Illumina TrueSeq amplification primers (Illumina, San Diego, California, USA). Cycle number was limited to 14 Cycles. Pooling and cleanup of Genotyping by Sequencing (GBS) libraries was done using 5 μl from each of the 96 amplified libraries. PCR primer and small amplicons were removed by Agencourt XP bead purification (Beckman Coulter Life Sciences, Indianapolis, USA) using 1 volume of beads. The PCR enzyme was removed by an additional purification on Qiagen MinElute Columns (Qiagen, Marylan, USA). The pooled Library was eluted in a final volume of 20μl Tris Buffer (5 mM Tris/HCl pH: 9).

Normalization was done using Trimmer Kit (Evrogen, Moscow, Russia). One μg pooled GBS library in 12 μl was mixed with a 4 μl 4x hybridization buffer, denatured for 3 min at 98°C and incubated for 5 hours at 68°C to allow reassociation of DNA fragments. Twenty μl of 2 x DSN master buffers was added, and the samples were incubated for 10 min at 68°C. One unit of DSN enzyme (1U/μl) was added and the reaction was incubated for another 30 min. Reaction was terminated by the addition of 20μl DSN Stop Solution, purified on a Qiagen MinElute Column (Qiagen, Marylan, USA) and eluted in 10μl Tris Buffer (5 mM Tris/HCl pH: 9). The normalized library pools were re-amplified in 100μl PCR reactions using MyTaq (Meridian Bioline, Memphis, Tennessee, USA). For each pool, a different i5-Adaptor primer was used to include i5-Indices into the libraries, allowing parallel sequencing of multiple libraries on the Illumina NextSeq 500 sequencer (Illumina, San Diego, California, USA). Cycle number was limited to 14 cycles. The GBS libraries were size selected on Blue Pippin (Sage Science, Massachusetts, USA), followed by a second size selection on a UltraPure™ Low Melting Point Agarose (LMP-Agarose gel; Thermo Fisher Scientific, Massachusetts, USA), removing fragments smaller than 300bp and those larger than 400bp. Sequencing was done on an Illumina NextSeq 500 (Illumina, San Diego, California, USA) using V2 Chemistry (300 cycles).

### Data processing

For data preprocessing restriction enzyme site filtering of read 5’ ends was performed and reads with 5’ ends not matching the restriction enzyme site were discarded. Quality trimming of adapter clipped Illumina reads was done by removing reads containing Ns, and trimming reads at 3’-end to get a minimum average Phred quality score of 20 over a window of ten bases. Reads with final length < 20 bases were discarded. A subsampling (evenly across the complete FASTQ files) of quality-trimmed reads was performed to 1.5 million read pairs per sample. FastQC reports (Andrews, 2008) were prepared for all FASTQ files and read counts were recorded.

Two software were tested to call variants for the pilot study with the goal of finding the best performing program (de-novo or using a reference genome): DiscoSnpRad (Gauthier et al., 2017), and Stacks (Catchen et al., 2013). The best performing restriction enzyme and best bioinformatics pipelines to call variants were subsequently used for this study (Table S1). All pipelines used here can be found at https://github.com/TheAriasLab/POPULATIONS_GENOMICS

For Stacks, Bowtie2 (Langmead & Salzberg, 2012) was implemented to index the “Hainan Tall Coconut” draft reference genome (Xiao et al., 2017) and to align reads to the reference genome index. Samtools (Li et al., 2009) was used to compress SAM files and to convert them to their binary version (BAM), necessary for Stacks. The Stacks script “*populations*” was implemented to filter and identify Single Nucleotide Polymorphisms (SNPs) according to: a minimum percentage of individuals (80) in a sampled site in which a SNP must be present to be considered, a minimum allele frequency (0.05) of a SNP to be considered (Fig. S2), and a maximum observed heterozygosity (0.70) of a SNP to be considered.

For DiscoSnpRad, the script “*run_discoSnpRad.sh*” was ran with the following parameters: remove low complexity bubbles, extend each polymorphism with left and right unitigs, k-mer size = 31, branching strategy, allows non-symmetrical branching bubbles = 1, minimum coverage = 12, maximum errors per read = 8, maximum deletion size = 10, maximum SNPs per bubble = 5. Other parameters were left as default. DiscoSnpRad SNPs were filtered according to the Stacks “*populations*” script.

Stacks took longer to run than DiscosnpRad and used more RAM but recovered more SNPs with less missing data (56 - 8.57%; Table. S1). The data set resulting from Stacks contains 27600 SNPs for 125 individuals, after removing accessions 3527 and 3418 (Table S1) due to 100% missing data rate.

### Population statistics, population structure and clustering analysis

R package Hierfstats 0.04-30 (Goudet, 2005) was used to calculate pairwise FST using Nei’s coefficients, inbreeding coefficients FIS and heterozygosity Ho. Confidence intervals of 95% in each sampled site, FIS and pairwise FST were estimated with 1000 bootstraps. Also, pairwise FST, inbreeding coefficients FIS and expected and observed heterozygosity (He, Ho) were calculated between the Pacific and Atlantic groups using “*basic.stats*” function of Hierfstat 0.04-30 R package (Goudet, 2005). Adegenet 2.1.1 (Jombart, 2008) was used initially to visualize genetic differences between sampled sites using Discriminant Analysis of Principal Components (DAPC). Poppr (Kamvar et al., 2015) was used to calculate Euclidean distances between individuals and to build a Minimum Spanning Network (MSN) to visualize genetic groupings. To visualize genetic differences FIS values calculated per accession in VCFtools (Danecek et al. 2011), and associate with phenotypic characteristics (see plant material and experimental design section) using PCA analysis in R.

### Linkage disequilibrium analysis

The extent of LD decay was assessed for the whole collection. Pairwise r2 values were estimated using the --geno-r2 function implemented in VCFtools for unphased genotypes (Danecek et al., 2011). To determine the decay of LD with increasing distance between loci (SNPs), the average r2 was expressed as a function of distance between SNPs (up to 1500Kb). A second round of LD calculations were done using the squared correlation coefficient for subgroups of SNPs from accessions of the Atlantic and Pacific coasts of Colombia independently.

### Genetic structure of populations

A Bayesian method Structure 2.3.4 (Hubisz et al., 2009) used to identify genetic structure of populations with an admixture model, a burn-in of 25,000, and a number of MCMC replicates after burn-in of 250,000 was run for the dataset (125 individuals and 27.600 SNPs). Runs were performed from k = 1 to k = 6, assuming coconut populations from Colombia could show population substructure based on a global divide between Atlantic and Pacific populations (Gunn et al., 2011) and further substructures within each coast. Each run was repeated 10 times for a total of 60 runs per test. Evanno’s method (Evanno et al., 2005), in house R scripts were used to calculate delta K as a measure that best describes the number of clusters in the data. Permutation of the runs were done in Clumpp 1.1.2 (Jakobsson & Rosenberg, 2007) and visualization of the data with inhouse R scripts. Hybrids were identified and filtered for some analysis using snapclust function in adegenet (Jombart, 2008). To visualize the Structure 2.3.4 results spatially we performed spatial interpolation of ancestry coefficients using “*maps*” function from the “*POPSutilities.R*” suite of functions (Jay et al. 2012).

A further look at population structure was inferred based on nearest neighbor haplotype “coancestry” implemented in RADpainter and fineRADstructure (Malinsky et al., 2018). For this analysis we used a set of filtered SNPs that are on Linkage Equilibrium.

### Phylogenetic networks

Phylogenetic networks represent conflicting signals in non-treelike processes, such as hybridization followed by introgression. The network displays relative evolutionary distances between taxa as well as uncertainty in the groupings in the form of “splits “of internal branches. To understand relationships and distances between samples, we used SplitsTree4 (Huson & Bryant, 2005) to infer a phylogenetic network with the Neighbor-net algorithm.

## Declarations

### Availability of data and materials

All data have been deposited in Bioproject (PRJNA579494), SRA (SRR10345275-SRR10345401) agronomic data (https://doi.org/10.6084/m9.figshare.13020623), morphology data (https://doi.org/10.6084/m9.figshare.12783470) and LD data (https://doi.org/10.6084/m9.figshare.13020635)

### Author contributions

JMMP: Co-developed questions and framework, performed analyses, wrote text.

GPC: Co-developed questions, obtained funding, made collections, and performed lab work.

LL: Co-developed questions, obtained funding, made collections, and performed lab work.

TAG: Co-developed questions and framework, performed analyses, mentored student author, assisted with obtaining funding, wrote and edited text.

## Acknowledgments

We thank Juan Carlos Bedoya from El Colegio Mayor de Antioquia, David Granada and Sara Ramirez from La Corporación para Investigaciones Biológicas (CIB) for help with fieldwork. Colin Findley, Diego Mauricio Riaño-Pachón, Andrew Crawford, Craig Barrett, Nhora Helena Ospina-Calderon, Alejandro Zuluaga, Camilo Chacon-Duque, and Ivan Soto-Calderón for comments on the paper. Luis Eduardo Mejia for help with figures and Juan David Pineda Cardenas for advice about computational resources used through EAFIT. This work was funded through the Colombian Research Grants Administration System (Ministerio de Ciencia y Tecnología) to GPC and TA 221371353189.

