## Supplementary information for "Genome-wide diversity of coconut from northern South America uncovers genotypes present in Colombia and strong population structure"

SUPPORTING INFORMATION

**Figure S1.** Comparison of two pipelines implemented to call SNPs. (a) Total number of SNPs filtered by its occurrence on at least one population (P1) to all populations (P5). (b) Approximately time needed to run (time), cores and RAM memory used to run pipelines (Cores/RAM), number of SNPs called for only one population and for SNPs called for all populations (SNPs), number of insertions, deletions or additions (Indels), average missing genotypes for individual basis for SNPs called in one population and for SNPs called for all populations (Missing Data), strategy used to identify SNPs, all analyses were carried out on Linux machines.

a.

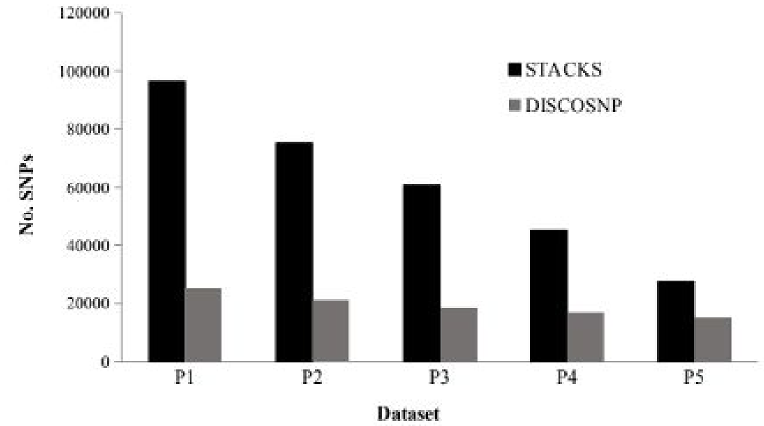


b.

| Pipeline | Time (h) | Cores/RAM (GB) | SNPs | Indels | Missing Data (%) | Method |
|---|---|---|---|---|---|---|
| Stacks | ~ 8 | 4/8GB | 96480 - 27600 | - | 56.36 – 8.57% | Reference genome |
| DiscoSnpRad | ~ 5 | 32/32GB | 25041 - 15435 | 1252 | 52.79 - 51.35 | *de novo* |

**Figure S2.** Minor Allele Frequency per loci histogram.

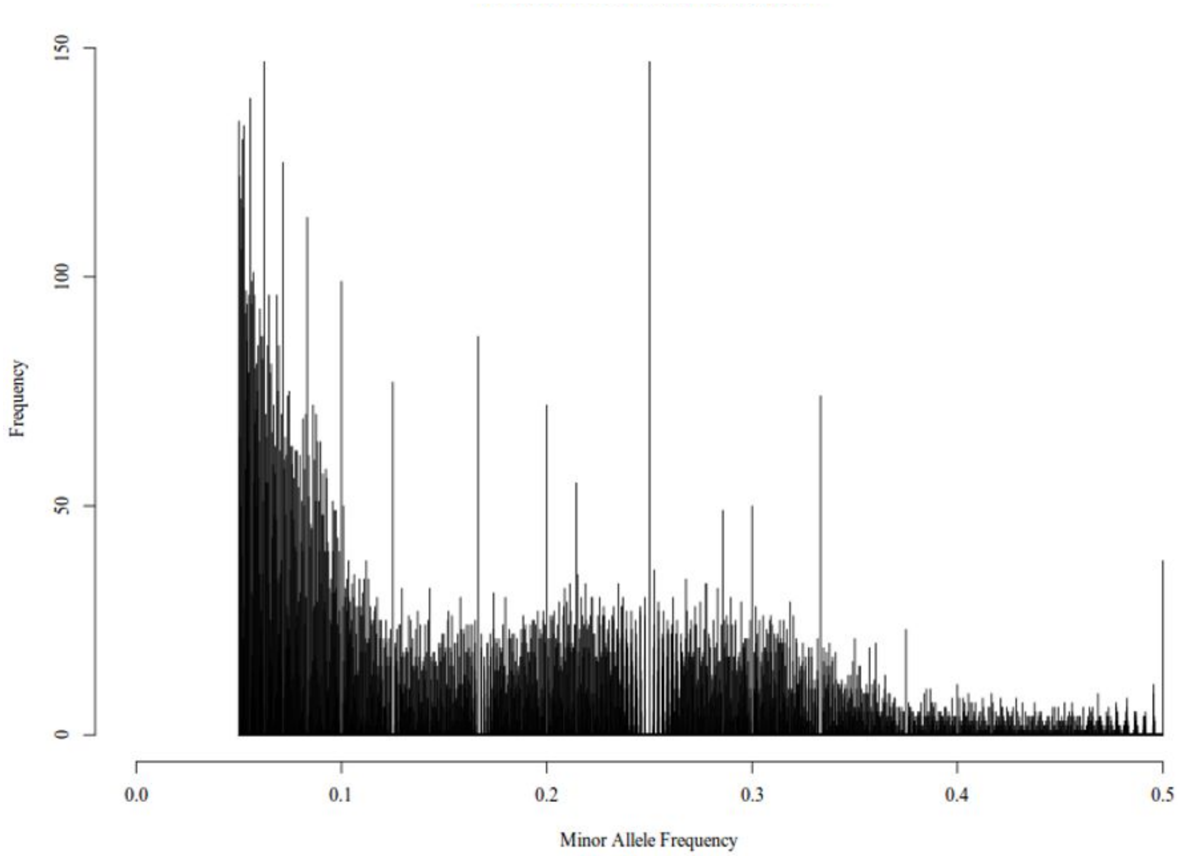


**Figure S3.** Population statistics per region and sample site. (a) Observed heterozygosity per region. (b) Expected heterozygosity per region. (c) Inbreeding coefficient per sample site. Boxes represent interquartile range, bars represent maximum and minimum values and the points represent potential outliers.

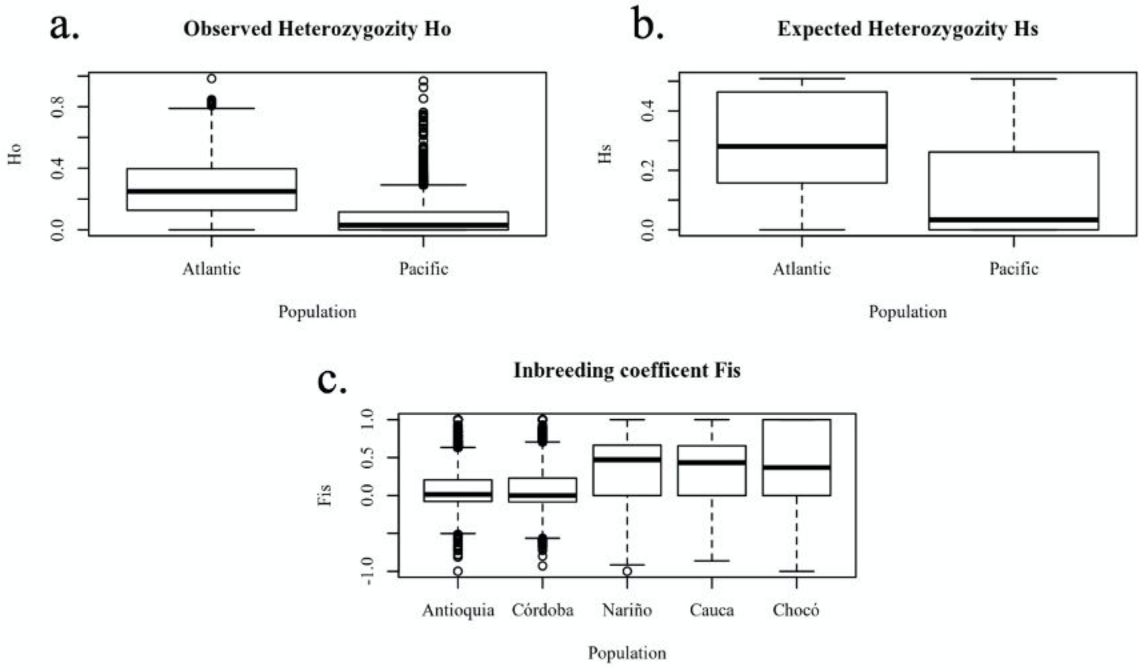

**Figure S4.** Population structure of Northern South America cultivated coconuts using SNPs recovered by STACKS and calculated in the larger dataset (125 accessions and 27600 SNPs). Delta K using all identified SNPs as markers.

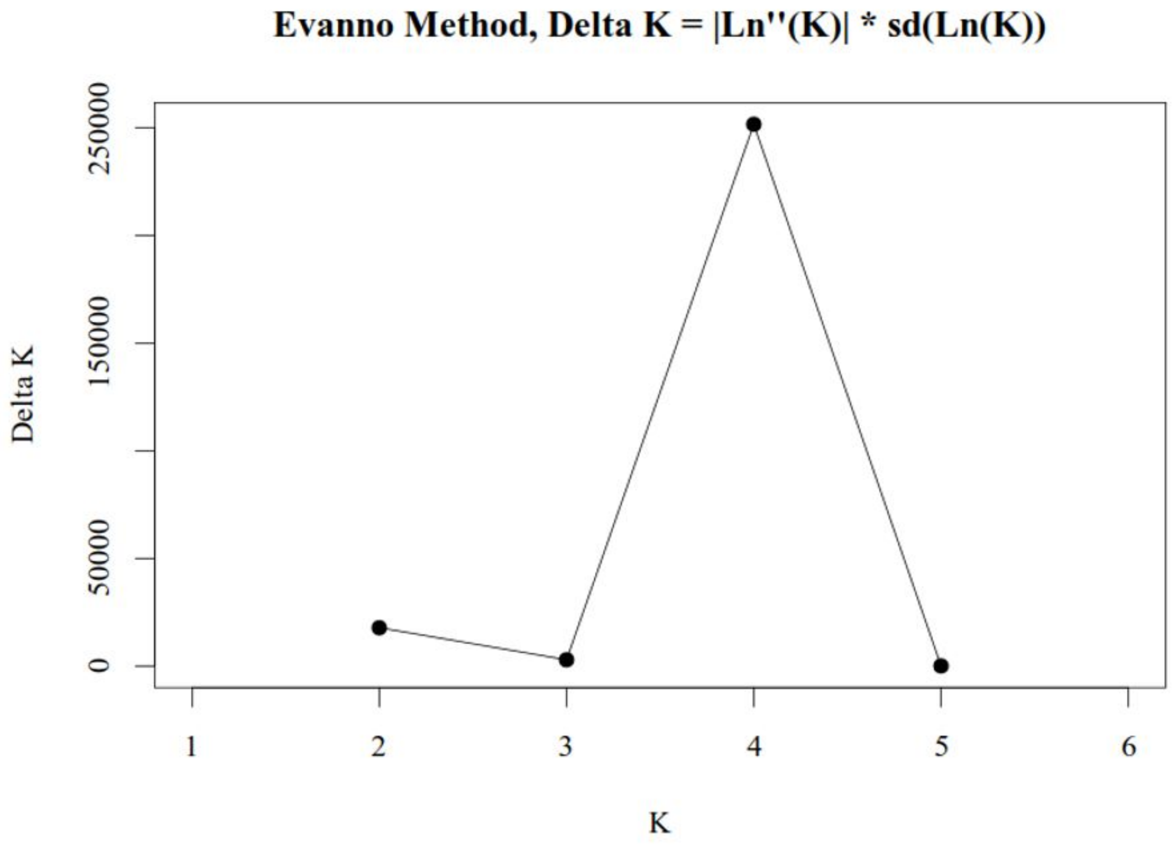


**Figure S5.** Population structure analysis of Northern South America cultivated coconuts using SNPs recovered by STACKS calculated in the larger dataset (125 accessions and 27600 SNPs)

and using *STRUCTURE.* A vertical bar represents each accession. Each color represents one ancestral population, and the length of each colored segment in each vertical bar represents the proportion contributed by ancestral populations. (a) *STRUCTURE* analysis for K = 2. (b) *STRUCTURE* analysis for K = 3.

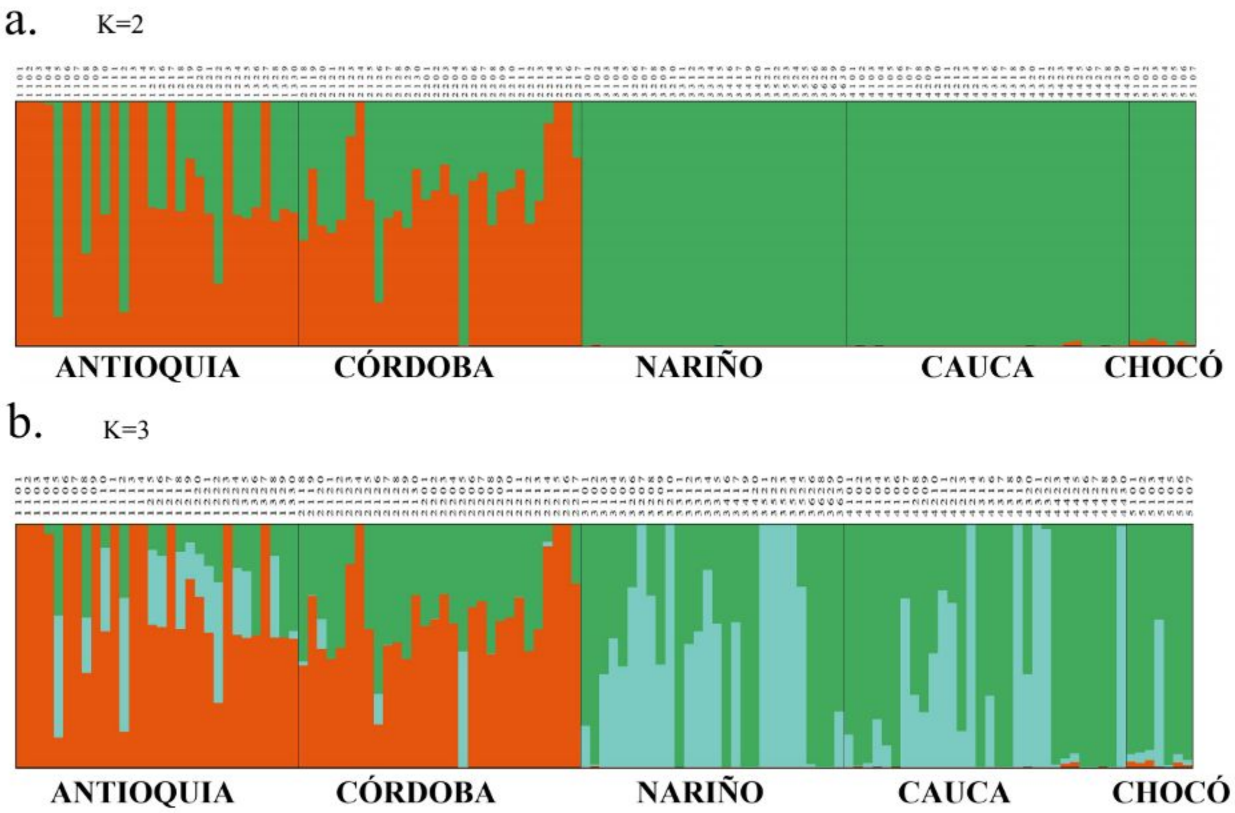


**Figure S6.** Population structure analysis of northern South America cultivated coconuts using *STRUCTURE.* a) dataset filtering all SNPs in Linkage Disequilibrium (LD: r2 > 0.3) (13900). A vertical bar represents each accession. Each color represents one ancestral population, and the length of each colored segment in each vertical bar represents the proportion contributed by ancestral populations. Evanno analysis K = 2 and *STRUCTURE* analysis for K = 2. b) dataset filtering all recent hybrids.

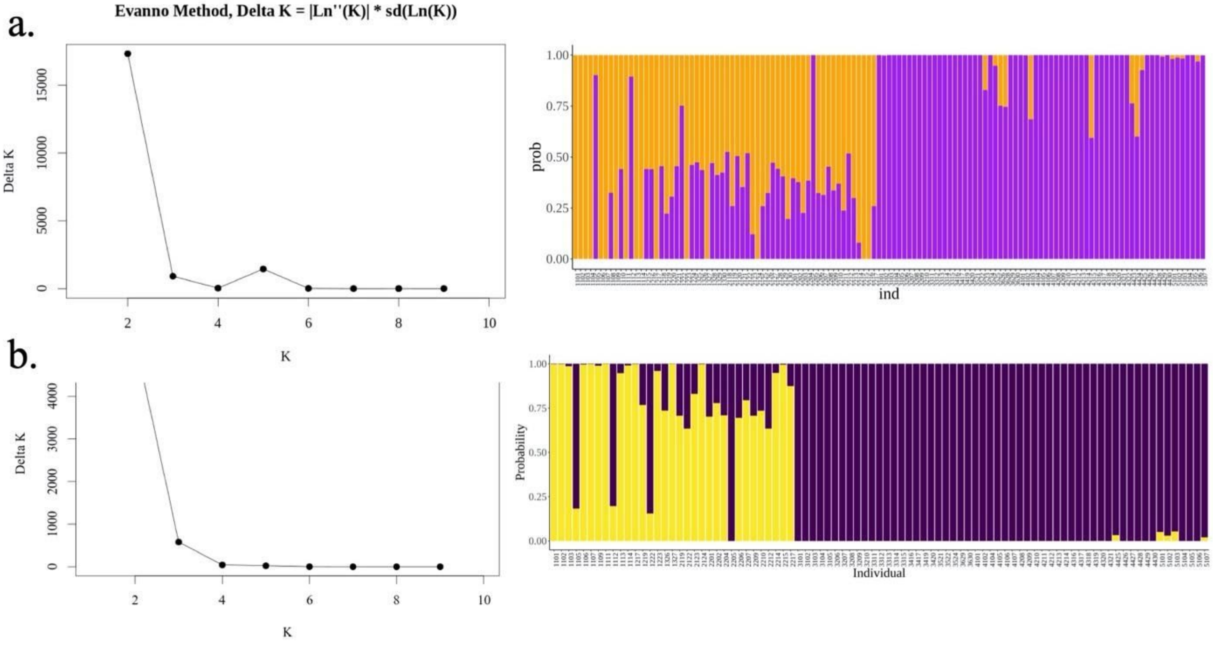


**Figure S7.** Coancestry matrix of Northern South America cultivated coconuts using SNPs recovered by STACKS and calculated using fineRADstructure in the larger dataset 27600 SNPs based on SNPs. The heat map depicts the high-resolution relationships of individuals selected across the Atlantic and Pacific Oceans of Colombia.

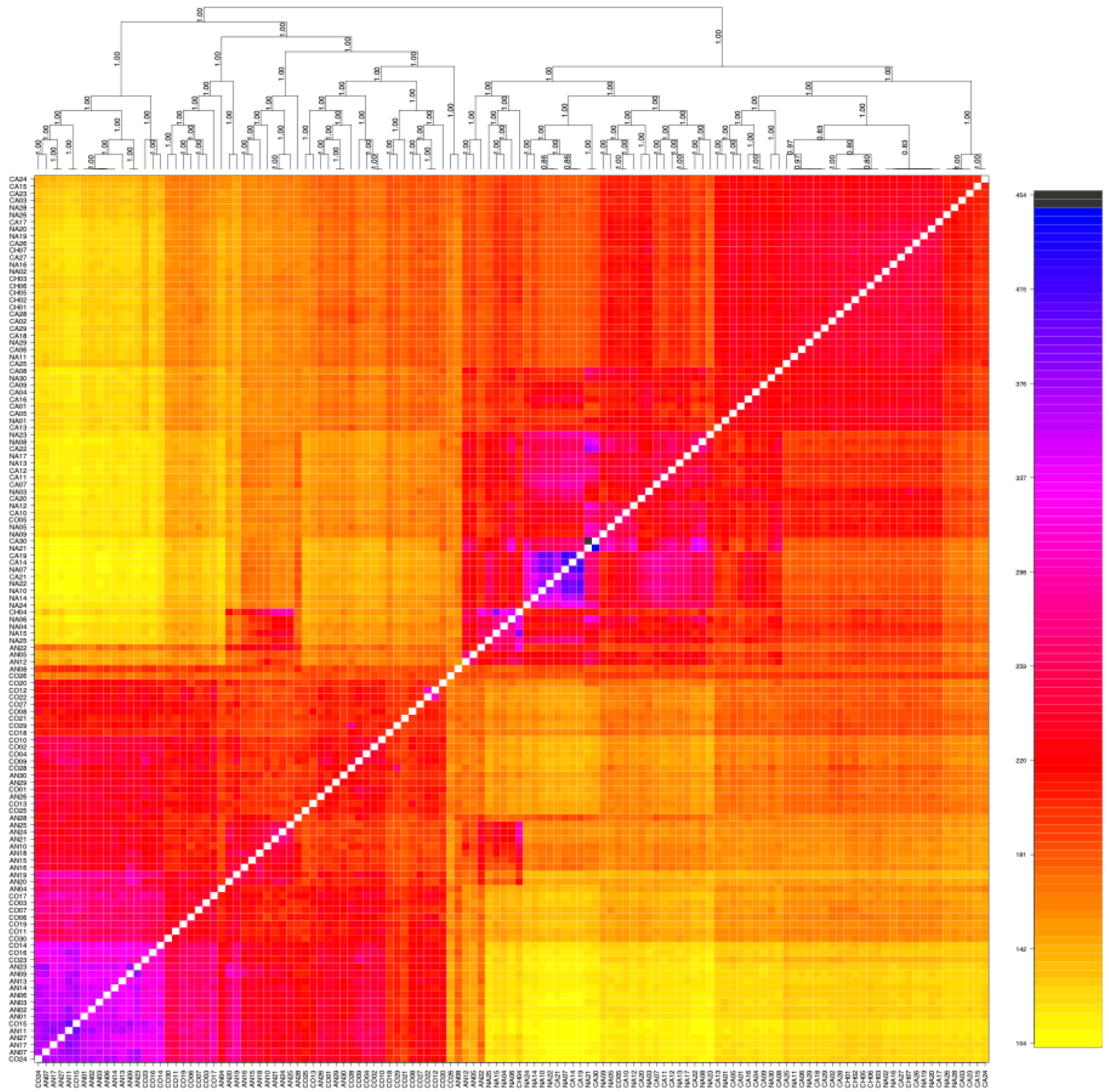

**Figure S8.** Principal components analysis for morphological and agronomic data using mean values for measurements. a. Five morphometric characters of the palms (rachis length, plant height, number of leaflets per plant, stem circumference and the ratio between the equatorial and polar diameter for the nuts (Xnut)) measure in all accessions. b. PCA of agronomic data for three harvest seasons 2017-2019 including nut equatorial circumference ECN (cm), nut weight NW (g), copra weight CW (g), fiber weight FW (g), and volume of coconut water W (ml).

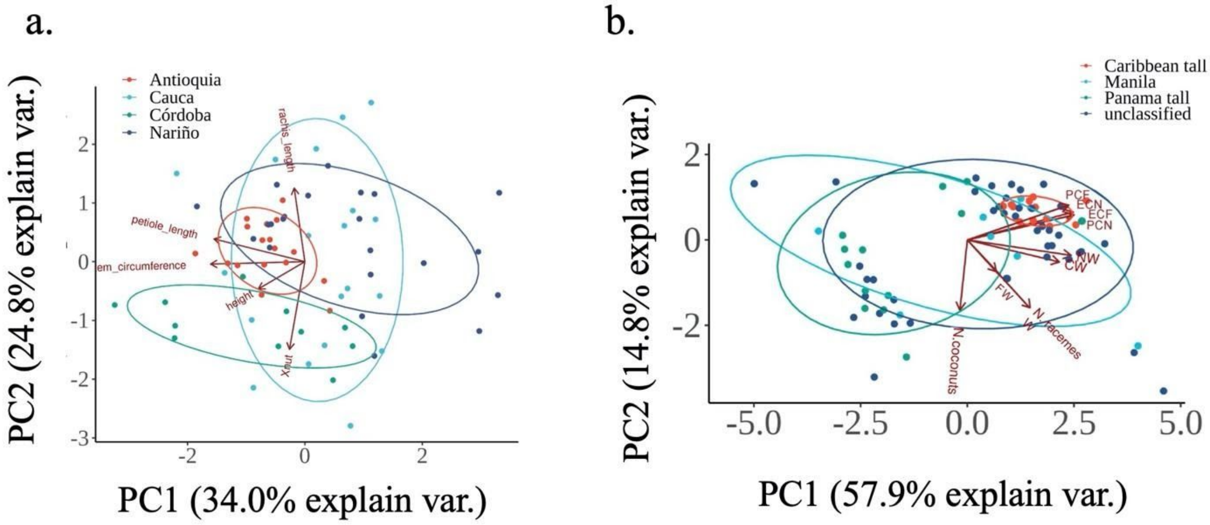


**Table S1.** Voucher, geographic coordinates and source information when available for northern South America coconut accessions.

| Locality | Code | Longitude | Latitude | Varieties |
|---|---|---|---|---|

| | | | | |
|---|---|---|---|---|
| **Caribbean Coast** | | | | |
| **Antioquia** | 1101 | -76.973 | 8.126575 | Alto Caribe |
| | 1102 | -76.9719916666667 | 8.12626944444444 | Alto Caribe |
| | 1103 | -77.0977388888889 | 8.12656388888889 | Alto Caribe |
| | 1104 | -76.9739277777778 | 8.12635 | Alto Caribe |
| | 1105 | -76.9735027777778 | 8.12615833333333 | NA |
| | 1106 | -76.9738888888889 | 8.12581111111111 | Alto Caribe |
| | 1107 | -76.9738944444444 | 8.12533055555556 | Alto Caribe |
| | 1108 | -76.9750805555556 | 8.12624444444445 | NA |
| | 1109 | -76.9750861111111 | 8.12443611111111 | NA |
| | 1110 | -76.9745805555556 | 8.12528333333333 | NA |
| | 1111 | -76.9743555555556 | 8.12605555555556 | Alto Caribe |
| | 1112 | -76.9722222222222 | 8.12479166666667 | NA |
| | 1113 | -76.9725083333333 | 8.12688611111111 | Alto Caribe |
| | 1114 | -76.9725388888889 | 8.12625833333333 | Alto Caribe |
| | 1215 | -76.9637027777778 | 8.12292222222222 | NA |
| | 1216 | -76.9643222222222 | 8.12268055555556 | NA |
| | 1217 | -76.9643166666667 | 8.12213888888889 | Alto Caribe |
| | 1218 | -76.9642861111111 | 8.12176388888889 | NA |

| | | | | |
|---|---|---|---|---|
| | 1219 | -76.9636666666667 | 8.12155833333333 | NA |
| | 1220 | -76.9633333333333 | 8.12177777777778 | NA |
| | 1221 | -76.9634 | 8.12202222222222 | NA |
| | 1222 | -76.9633361111111 | 8.12129722222222 | NA |
| | 1223 | -76.9636222222222 | 8.123175 | Alto Caribe |
| | 1224 | -76.9628472222222 | 8.12279722222222 | NA |
| | 1326 | -76.7418861111111 | 8.13921666666667 | NA |
| | 1327 | -76.7413972222222 | 8.13957777777778 | NA |
| | 1328 | -76.7417361111111 | 8.13983055555556 | Alto Caribe |
| | 1329 | -76.7414694444444 | 8.14045833333333 | NA |
| **Córdoba** | 2118 | -76.1930666666667 | 9.04446944444445 | NA |
| | 2119 | -76.1927444444445 | 9.043625 | NA |
| | 2120 | -76.1924111111111 | 9.04355277777778 | NA |
| | 2121 | -76.1934222222222 | 9.04265833333333 | NA |
| | 2122 | -76.1932972222222 | 9.04488333333333 | NA |
| | 2123 | -76.1835333333333 | 9.04438055555556 | NA |
| | 2124 | -75.9680555555556 | 9.05265555555556 | NA |
| | 2125 | -76.2177694444444 | 9.05320277777778 | Alto Caribe |
| | 2126 | -76.2669277777778 | 9.05323333333333 | NA |

| | | | |
|---|---|---|---|
| 2128 | -76.1683416666667 | 9.0472 | NA |
| 2129 | -76.1693416666667 | 9.04626388888889 | NA |
| 2130 | -76.1684388888889 | 9.04576111111111 | NA |
| 2201 | -76.2252055555555 | 9.06939166666667 | NA |
| 2202 | -76.2252055555555 | 9.06939166666667 | NA |
| 2203 | -76.2258805555556 | 9.07429722222222 | NA |
| 2204 | -76.2260416666667 | 9.06928611111111 | NA |
| 2205 | -76.2257694444444 | 9.07549444444445 | NA |
| 2206 | -76.2226166666667 | 9.07441388888889 | NA |
| 2207 | -76.2221083333333 | 9.07361944444445 | NA |
| 2209 | -76.22275 | 9.07770277777778 | NA |
| 2210 | -76.2234527777778 | 9.07728611111111 | NA |
| 2211 | -76.4316388888889 | 8.16589444444445 | NA |
| 2212 | -76.4330472222222 | 8.16532222222222 | NA |
| 2213 | -76.4330472222222 | 8.166725 | NA |
| 2214 | -76.4327694444444 | 8.16667777777778 | NA |
| 2215 | -76.432725 | 8.16531388888889 | Alto Caribe |
| 2216 | -76.4715416666667 | 8.16370833333333 | Alto Caribe |
| 2217 | -76.47145 | 8.1638 | NA |

| | | | | |
|---|---|---|---|---|
| **Pacific Coast** | 3101 | -78.9438083333333 | 1.117375 | NA |
| **Nariño** | 3102 | -78.9437944444444 | 1.117525 | NA |
| | 3103 | -78.9438444444444 | 1.11718888888889 | NA |
| | 3104 | -78.9438972222222 | 1.11704166666667 | NA |
| | 3105 | -78.9443972222222 | 1.11688611111111 | Alto Pacífico |
| | 3206 | -78.8926527777778 | 1.12568333333333 | NA |
| | 3207 | -78.8927583333333 | 1.12587222222222 | Manila |
| | 3209 | -78.8932361111111 | 1.12506944444444 | NA |
| | 3210 | -78.892 | 1.12480833333333 | Manila |
| | 3311 | -78.8902611111111 | 1.13105277777778 | Alto Pacífico |
| | 3312 | -78.8907444444445 | 1.13229722222222 | NA |
| | 3313 | -78.89045 | 1.13286944444444 | NA |
| | 3314 | -78.8899861111111 | 1.13161666666667 | NA |
| | 3315 | -78.8901027777778 | 1.13113888888889 | NA |
| | 3416 | -78.8907777777778 | 1.12591388888889 | NA |
| | 3417 | -78.8902361111111 | 1.126975 | NA |
| | 3419 | -78.890625 | 1.12610833333333 | Alto Pacífico |
| | 3420 | -78.890625 | 1.12599722222222 | Alto Pacífico |
| | 3521 | -79.0505555555556 | 1.11813611111111 | Manila |

| | | | | |
|---|---|---|---|---|
| | 3522 | -79.0444055555556 | 1.118575 | Manila |
| | 3523 | -79.0450472222222 | 1.11881111111111 | Manila |
| | 3524 | -79.04585 | 1.1182 | Manila |
| | 3525 | -79.04595 | 1.11825277777778 | NA |
| | 3626 | -78.9942694444444 | 1.11444166666667 | Alto Pacífico |
| | 3628 | -78.9958333333333 | 1.11449166666667 | Alto Pacífico |
| | 3629 | -78.9941972222222 | 1.11511388888889 | Alto Pacífico |
| | 3630 | -78.9942861111111 | 1.11497222222222 | Injerto verde |
| **Cauca** | 4101 | -78.3992888888889 | 2.10985833333333 | NA |
| | 4102 | -78.3993861111111 | 2.10975833333333 | Alto Pacífico |
| | 4103 | -78.3996138888889 | 2.11047777777778 | Alto Pacífico |
| | 4104 | -78.39995 | 2.11054166666667 | NA |
| | 4105 | -78.3999055555556 | 2.11129166666667 | NA |
| | 4106 | -78.3982222222222 | 2.11039444444444 | Alto Pacífico |
| | 4107 | -78.3984055555556 | 2.11135555555556 | NA |
| | 4209 | -78.4350055555556 | 2.10880277777778 | NA |
| | 4210 | -78.4346222222222 | 2.10837222222222 | NA |
| | 4211 | -78.4354805555556 | 2.11044166666667 | NA |
| | 4212 | -78.4353805555556 | 2.10982777777778 | NA |

| | | | | |
|---|---|---|---|---|
| | 4213 | -78.4354138888889 | 2.10997222222222 | NA |
| | 4214 | -78.4343972222222 | 2.10825277777778 | Manila |
| | 4316 | -78.5326666666667 | 2.107175 | NA |
| | 4317 | -78.5326166666667 | 2.10706666666667 | Alto Pacífico |
| | 4318 | -78.5322861111111 | 2.107 | Alto Pacífico |
| | 4319 | -78.5301138888889 | 2.10516944444444 | Manila |
| | 4320 | -78.52985 | 2.10503888888889 | NA |
| | 4321 | -78.5304722222222 | 2.10517222222222 | Manila |
| | 4322 | -78.5309138888889 | 2.10721111111111 | Manila |
| | 4423 | -78.5119194444444 | 2.113525 | Alto pacífico |
| | 4424 | -78.5121555555556 | 2.11420277777778 | Alto pacífico |
| | 4425 | -78.51185 | 2.11445555555556 | Alto pacífico |
| | 4428 | -78.5108027777778 | 2.11384722222222 | Alto pacífico |
| | 4429 | -78.5098111111111 | 2.11396388888889 | Alto Pacífico |
| | 4430 | -78.5104416666667 | 2.11301666666667 | Manila |
| **Choco** | 5101 | -77.3511638888889 | 5.60071388888889 | Alto Pacífico |
| | 5102 | -77.3512055555556 | 5.6008 | Alto Pacífico |
| | 5103 | -77.3512333333333 | 5.60082222222222 | Alto Pacífico |
| | 5104 | -77.351125 | 5.60078611111111 | NA |

| | | | |
|---|---|---|---|
| 5105 | -77.3509638888889 | 5.60086666666667 | Alto Pacífico |
| 5106 | -77.3506 | 5.601 | Alto Pacífico |
| 5107 | -77.3505361111111 | 5.60103611111111 | Alto Pacífico |

**Table S2.** Phenotypic traits in Coconut accessions cultivated in Colombia, number of racemes (NR), Number of coconuts (NC), Polar circumference of the fruit (PCF, cm), equatorial circumference of the fruit (ECF, cm), polar circumference of the nut (PCN, cm), equatorial circumference of the nut (ECF, cm), copra weight (CW, g), and Fiber weight (FW, g) and water inside the fruit (W, ml)

| Code | Group | NR | NC | PCF | ECF | PCN | ECN | NW | CW | FW | Water |
|---|---|---|---|---|---|---|---|---|---|---|---|
| 1101 | Atlantic tall | 1 | 7 | 57.67 | 52.33 | 34.33 | 34 | 576.67 | 277.67 | 648 | 121.67 |
| 1102 | Atlantic tall | 1 | 8.33 | 59.33 | 45.67 | 33.67 | 31.33 | 519 | 240.33 | 712.33 | 106.67 |
| 1103 | Atlantic tall | 2 | 11.33 | 53 | 51.33 | 34 | 33.5 | 522.33 | 263 | 630.33 | 85 |
| 1105 | hybrid | 1 | 14 | 56 | 43 | 33.67 | 30.33 | 442 | 259.67 | 503.33 | 85 |

| | | | | | | | | | | | |
|---|---|---|---|---|---|---|---|---|---|---|---|
| 1106 | Atlantic tall | 1 | 7 | 62.67 | 49.67 | 36 | 32 | 655.67 | 310.67 | 620.67 | 206.67 |
| 1107 | Atlantic tall | 1 | 6 | 58.67 | 48.33 | 37 | 33 | 636.67 | 292.3 | 654.67 | 110 |
| 1109 | hybrid | 2 | 13 | 56.33 | 48.5 | 33.5 | 31.17 | 513 | 203.37 | 446.33 | 121.67 |
| 1111 | Atlantic tall | 1.6 | 11.66 | 57.67 | 49.17 | 33.33 | 32.67 | 573.337 | 222.67 | 452.337 | 165.67 |
| 1112 | hybrid | 1 | 8.67 | 57 | 50.33 | 33 | 31 | 537 | 190.3 | 435.337 | 126.67 |
| 1113 | Atlantic tall | 1 | 9 | 60.37 | 53.33 | 34.67 | 32.67 | 581.33 | 268.33 | 635.67 | 130 |
| 1114 | Atlantic tall | 1 | 10 | 60.67 | 57.67 | 34 | 31.5 | 622 | 293.67 | 568.67 | 121.67 |
| 1217 | Atlantic tall | 1 | 11.67 | 57 | 51.67 | 32.5 | 29.75 | 483.33 | 230.67 | 693.67 | 80 |
| 1219 | hybrid | 1.6 | 10 | 58.33 | 44 | 34 | 30.67 | 554.67 | 274.33 | 454.67 | 148.33 |
| 1222 | hybrid | 1 | 8 | 56.5 | 50.5 | 34.33 | 31 | 547 | 253.67 | 483.33 | 153.33 |
| 1223 | Atlantic tall | 1 | 7.67 | 60 | 48.33 | 39 | 36 | 606.67 | 276.33 | 619 | 191.67 |
| 1326 | hybrid | 1 | 14.67 | 62.25 | 50 | 36.17 | 37.5 | 679.33 | 300.67 | 607.67 | 150 |
| 1327 | Atlantic tall | 1 | 7.33 | 77.33 | 61 | 40 | 37.5 | 701.67 | 368.67 | 1068 | 100 |
| 2119 | hybrid | 9.8 | 11.14 | 56.11 | 49.41 | 32.97 | 34.31 | 480.71 | 454 | 170.43 | 164.71 |
| 2122 | hybrid | 9.7 | 13.28 | 56.95 | 51.62 | 33.7 | 34.58 | 523.36 | 504.95 | 168.43 | 190 |
| 2123 | hybrid | 8.5 | 9.83 | 57.56 | 49.74 | 34.33 | 35.32 | 475 | 451.75 | 176.75 | 180.67 |
| 2124 | Atlantic tall | 7.43 | 8 | 55.5 | 45.67 | 33.5 | 34.75 | 476.43 | 440 | 173.6 | 177.14 |
| 2201 | hybrid | 11 | 19.67 | 62 | 52.1 | 33.73 | 33.07 | 518.47 | 473.33 | 41.67 | 180 |
| 2202 | hybrid | 10 | 17.5 | 62.6 | 52.8 | 34.475 | 33.45 | 514.45 | 468.25 | 41.5 | 177.5 |
| 2206 | hybrid | 8.5 | 10 | 60.23 | 56.13 | 32.4 | 31.775 | 574.45 | 525.6 | 41 | 211.5 |
| 2207 | hybrid | 10 | 17.75 | 61.67 | 58.43 | 33.93 | 34.65 | 578.6 | 533.6 | 41.65 | 214 |
| 2209 | hybrid | 11 | 8.75 | 59.67 | 51.65 | 33.725 | 32.65 | 661.3 | 612.4 | 41.5 | 249 |

| | | | | | | | | | | | |
|---|---|---|---|---|---|---|---|---|---|---|---|
| 2210 | hybrid | 8.75 | 8.25 | 59.67 | 51.97 | 33.87 | 33.07 | 663.45 | 618 | 42.15 | 264.35 |
| 2212 | hybrid | 10.25 | 4.5 | 59.62 | 52.5 | 33.825 | 34.1 | 409.35 | 297.75 | 39.03 | 154.75 |
| 2215 | Atlantic tall | 8.75 | 3.5 | 56.85 | 52.2 | 33.3 | 32.85 | 675.75 | 643.2 | 40.95 | 212.25 |
| 2217 | hybrid | 8.75 | 5 | 56.67 | 47.83 | 32.8 | 32.225 | 485.75 | 445.75 | 39.225 | 176 |
| 3101 | hybrid | 1 | 4 | 58 | 49 | 37 | 35 | 749 | 323 | 440 | 110 |
| 3102 | hybrid | 1 | 5 | 51 | 41.5 | 33.5 | 28 | 418 | 190 | 375 | 86 |
| 3103 | hybrid | 2 | 4 | 51 | 45.5 | 33.5 | 33 | 580 | 297 | 344 | 106 |
| 3206 | hybrid | 49.76 | 2.86 | 41.86 | 53.75 | 40.54 | 34.86 | 882.57 | 409.47 | 554.35 | 360.49 |
| 3207 | Dwarf | 45.4 | 2.71 | 41.15 | 50.93 | 38.27 | 34.01 | 1101.79 | 425.19 | 578.29 | 330.09 |
| 3208 | hybrid | 46.19 | 4.14 | 40.37 | 51.76 | 39.58 | 34.20 | 912.14 | 649.17 | 607.64 | 445.36 |
| 3209 | hybrid | 29.91 | 12.43 | 40.24 | 39.96 | 34.02 | 29.65 | 459.55 | 216.17 | 332.96 | 160.97 |
| 3210 | Dwarf | 24.14 | 4 | 38.36 | 41.52 | 32.86 | 27.34 | 398.14 | 206.79 | 335.60 | 140.31 |
| 3311 | Panama tall | 1.8 | 3 | 58.7 | 55.77 | 40.15 | 39.46 | 681.87 | 372.9 | 760.38 | 224.5 |
| 3312 | hybrid | 1.8 | 3.2 | 58.71 | 55.67 | 37.66 | 38 | 735.73 | 263.12 | 550.47 | 207 |
| 3313 | hybrid | 1.8 | 5 | 58.5 | 55.96 | 39.39 | 36.88 | 707.5 | 279.6 | 601.2 | 178 |
| 3314 | hybrid | 2 | 6.8 | 47.37 | 39.6 | 33.81 | 29.572 | 383 | 196.54 | 354.6 | 107.266 |
| 3315 | hybrid | 1.4 | 4.8 | 45.82 | 40.66 | 34.16 | 31.39 | 488.4 | 248.51 | 476.7 | 156.03 |
| 3416 | hybrid | 1.4 | 1.8 | 45.04 | 40.28 | 35.58 | 33.16 | 444.24 | 199.66 | 502.36 | 129.52 |
| 3417 | hybrid | 0.6 | 1.4 | 18.12 | 14.4 | 11.4 | 9.8 | 208.8 | 62.7 | 154.5 | 25.1 |
| 3419 | Panama tall | 1.2 | 2.8 | 33.66 | 29.9 | 29.54 | 26.98 | 309.6 | 232.9 | 234.2 | 103.2 |
| 3420 | Panama tall | 1.4 | 3.6 | 44.78 | 37.35 | 32.36 | 30.68 | 384.83 | 151.8 | 355.99 | 110.46 |
| 3521 | Dwarf | 2 | 14 | 51 | 40 | 28 | 25 | 635 | 467 | 730 | 120 |
| 3522 | Dwarf | 1 | 3 | 55.8 | 53 | 39 | 35.7 | 308.5 | 210 | 380 | 240 |
| 3524 | Dwarf | 1 | 11 | 52.5 | 41 | 30.5 | 26.5 | 470 | 215 | 482 | 125 |

| | | | | | | | | | | | |
|---|---|---|---|---|---|---|---|---|---|---|---|
| 3629 | Panama tall | 0.75 | 2.25 | 29.025 | 28.55 | 26.3 | 24.7 | 622.125 | 230.375 | 449.125 | 130.25 |
| 3630 | hybrid | 1.25 | 3 | 62.675 | 57.5 | 51.25 | 37.075 | 667.625 | 306.625 | 671.75 | 358.25 |
| 4101 | hybrid | 1.4 | 18.8 | 16.1 | 14.5 | 10.5 | 9.6 | 302.6 | 182.2 | 551 | 197 |
| 4102 | Panama tall | 1.6 | 8.6 | 16.3 | 12.6 | 11.6 | 9.2 | 261.2 | 169.6 | 388.6 | 115 |
| 4104 | hybrid | 2.2 | 16.8 | 16.2 | 15.2 | 10.25 | 11.2 | 326.6 | 200.4 | 560.2 | 95 |
| 4105 | hybrid | 1.4 | 13.8 | 15.375 | 13.7 | 9.7 | 9.7 | 282.2 | 166.8 | 327.4 | 166.6 |
| 4106 | Panama tall | 2.2 | 22.2 | 16.6 | 15.1 | 11.1 | 10.2 | 383.4 | 236.6 | 746.6 | 268.2 |
| 4107 | hybrid | 1.4 | 10 | 14.875 | 11.8 | 9.2 | 10.4 | 260.4 | 115 | 407 | 158.2 |
| 4209 | hybrid | 2.6 | 16.2 | 17.9 | 16 | 12.64 | 10.7 | 479 | 209.4 | 631.6 | 235.6 |
| 4210 | hybrid | 3.4 | 38.8 | 14.6 | 12.7 | 11.06 | 10.9 | 293 | 171.2 | 535.2 | 187.8 |
| 4211 | hybrid | 3.8 | 30 | 15.9 | 12.9 | 11.9 | 9.78 | 201.6 | 138.6 | 327.2 | 119 |
| 4212 | hybrid | 1.6 | 13.2 | 18.24 | 15.5 | 11.4 | 10.3 | 419 | 158.4 | 531.2 | 217 |
| 4213 | hybrid | 2.2 | 19.4 | 19.1 | 15.1 | 12.7 | 9.1 | 380 | 179.6 | 459.6 | 239.4 |
| 4316 | hybrid | 2.2 | 15.2 | 16.5 | 11.9 | 8.76 | 8.44 | 242.4 | 162.8 | 607.4 | 163 |
| 4317 | Panama tall | 2.2 | 12.8 | 14.5 | 11.4 | 9.9 | 8.54 | 241 | 152.6 | 547.6 | 179.6 |
| 4318 | Panama tall | 2.2 | 17.6 | 14.1 | 13.5 | 8.3 | 8.9 | 224.4 | 159.2 | 818 | 148 |
| 4321 | Dwarf | 1 | 4.5 | 9.5 | 8 | 5 | 5.5 | 136 | 86 | 407.5 | 78 |
| 4425 | Panama tall | 1.6 | 14.2 | 16.6 | 13.6 | 10.9 | 10 | 343.2 | 260.6 | 390.8 | 193.6 |
| 4426 | Panama tall | 1 | 6.5 | 11.5 | 10 | 9 | 7.5 | 185.5 | 142.5 | 345 | 135 |
| 4427 | Panama tall | 1 | 5 | 12 | 9.5 | 10 | 7.5 | 200.5 | 135.5 | 431.5 | 92.5 |
| 4428 | Panama tall | 2 | 15.8 | 16.1 | 13.7 | 8.4 | 7.7 | 227.8 | 157 | 297.4 | 92 |

| | | | | | | | | | | | |
|---|---|---|---|---|---|---|---|---|---|---|---|
| 4429 | Panama tall | 2.25 | 15.25 | 14.5 | 15.25 | 9.55 | 9 | 315.25 | 213.5 | 376.75 | 233.25 |
| 4430 | Dwarf | 2.2 | 13.6 | 19.2 | 15.2 | 11.26 | 9.4 | 331.6 | 245.2 | 620.4 | 241.4 |
